# RNA Polymerase II subunits overexpressions induce genome instability and deregulate transcription

**DOI:** 10.1101/2024.11.20.622938

**Authors:** Martina Muste Sadurni, Megan Louise Jones, Lucas Pavlou, Mohamed Radi, Smaragda Kompocholi, Paolo Passaretti, Agnieszka Gambus, Lea H. Gregersen, Marco Saponaro

## Abstract

Independently of the pathways or circuits deregulated in cancer cells, these present altered transcription patterns, often also direct consequence of deregulation of transcription factors. In this sense, also the RNA Polymerase complexes responsible for transcription can be affected in cancers. We find that upregulations of RNA Polymerase II subunits, especially the largest ones, correlates with poor cancer patients’ outcome across a range of tumor types, presenting increased genome instability. Overexpressing the subunits RPB1, RPB3 and RPB4 in cells we find that these induce DNA damage. However, the mechanisms behind this increased genome instability are specific for each subunit, linked to the unique transcription alterations generated by the subunit overexpression. Importantly, we find significant overlap between the genes with more DNA damage in our cell line models and those more affected in cancers with subunit upregulation, indicating that upregulations could be responsible for some of the phenotypes present in these patients.

## Introduction

Deregulated transcription is a well-established hallmark of cancer cells, consequence of deregulated signaling pathways or altered expression of transcription factors^1–3^. However, in it has become evident that together with sequence specific DNA-binding factors, also the basal transcription machinery of general transcription factors and RNA Polymerase complexes can be deregulated in cancer cells^4^. For example, RNA Polymerase I (RNAPI) is induced by the oncogene MYC to support cell growth, making cells reliant on RNAPI activity and sensitizing them to RNAPI inhibitors^5^. Equally, RNA Polymerase III (RNAPIII) has been long known to be highly expressed in cancer tissues^6^. Regarding RNA Polymerase II (RNAPII), several publications have highlighted how single subunits present deregulated expression levels and/or are mutated in many cancer types (reviewed in ^7^). Nevertheless, it remains still unclear whether deregulations of the subunits in cancers are associated with specific oncogenic drivers, or whether they could directly affect patients’ outcome contributing to specific cancer phenotypes. In the RNAPII, 3 out of the 12 subunits form the active site^8,9^, with structural studies showing that the largest subunits are central for the interaction with several transcription-associated complexes^10–12^. However, recent data have highlighted how each subunit may have specific roles during transcription, as degron degradation of any of the 12 RNAPII subunits presented specific transcription defects, even though degron degradation of one subunit was sometimes associated with destabilization of multiple subunits together^13^. Therefore, to determine which deregulations of the RNAPII complex subunits correlated more with reduced cancer patients’ survival, and whether they shared specific phenotypes across different cancer types, we performed an analysis of all The Cancer Genome Atlas (TCGA) datasets available through cBioPortal. This analysis found that in particular upregulations of the four largest subunits of the complex (genes POLR2A-D/subunits RPB1-4) correlated with poor outcomes in a range of tumors, characterized by increased genome instability, and not linked to a specific oncogenic driver. We also found poor to no correlation between the expression levels of the subunits, indicating that cancer cells would be experiencing deregulated expression of one subunit at the time. We hypothesized that non-stochiometric expression of single subunits could compete with the RNAPII on chromatin for transcription factors, consequently deregulating RNAPII transcription. To test this hypothesis, we established doxycycline inducible cell line models able to overexpress three of the largest subunits of the complex, RPB1/POLR2A, RPB3/POLR2C and RPB4/POLR2D. Using these systems and a combination of cell biology and genome-wide approaches, we found that overexpression of each subunit induced increased transcription activity, increased DNA damage levels and altered C-terminal domain (CTD) phosphorylation patterns of the RNAPII. Importantly, the DNA damage was specifically linked to the deregulations of the CTD induced by the overexpression of the single subunit. Moreover, we showed that the transcription defects and genome instability induced by RPB3/POLR3C overexpression were mainly caused by defective recruitment of BRD4 to the RNAPII complex. Finally, we uncovered that the genes with increased DNA damage levels in cells when we overexpressed a subunit, often presented higher levels of genomic alterations particularly in cancer patients with subunits’ upregulations. All in all, this study has identified how deregulated expressions of the RNAPII subunits could contribute directly to some of the phenotypes found in the affected patients. Moreover, it highlighted how deregulation of subunits of the complex might affect the interchange of transcription factors with the RNAPII complex on chromatin because of altered CTD phosphorylation patterns. This would lead to defective transcription, important for the efficient transcription of genes but also for genome stability maintenance.

## Material and methods

### Cancer genomic databases analysis

For the analysis of RPB1-12/POLR2A-L deregulations we assessed data through cBioPortal^14^ (http://cbioportal.org) updated in September 2024. For the pan-cancer analyses, we selected all the 32 PanCancer Atlas studies, with a total of 10967 patients. For the specific cancer datasets, we analyzed the Breast Invasive Carcinoma (BIC), Brain Lower Grade Glioma (BLGG), Kidney Chromophope (KICH), Kidney Renal Clear Cell Carcinoma (KIRC), Liver Hepatocellular Carcinoma (LIHC), Acute Myeloid Leukemia (AML), and Uterine Corpus Endometrial Carcinoma (UCEC), selecting Firehose Legacy studies as these had the largest cohorts of samples. “Complete samples” were analyzed for all cancer types except UCEC, for which we selected “All samples” as the number of “Complete samples” was low. To identify patients with upregulations in the subunits, we selected “Putative copy-number alterations from GISTIC”, “mRNA expression z-Scores – relative to diploid samples” with a z-Score threshold of >2.0 as suggested by default. “Fraction Genome Altered” and “Mutation Count” levels in both “altered” and “unaltered” patients was analyzed. The list of the most deregulated cancer driver genes was obtained from IntOGen^15^ (www.intogen.org). For the genomic alteration analysis, copy number alterations and mutations were analyzed separately from the “Comparison/Survival-Genomic Alterations” tab on cBioPortal, plotting how frequently a specific gene is found genomically altered in cancer patients with upregulation of the RNAPII subunit towards control patients without upregulation in any of the four largest subunits.

### Establishment of cell culture models

All cell lines were cultured in complete DMEM: Dulbecco’s Modified Eagle’s Medium - high glucose (Sigma), supplemented with 0.01 % Pen/Strep (Gibco), 0.01 % Glutamine (Gibco) and 10 % Tet-free Fetal bovine serum (Sigma) (FBS) or Tet-free FBS (TakaraBio), under selection of 5 μg/mL Blasticidin (ThermoFisher) and the appropriate selection marker in a humidified incubator at 37C and 5 % CO_2_. Unless otherwise stated, for the single overexpression of the RNAPII subunits cells were seeded at 20% confluence and induced with 5-10 ng/mL Doxycycline (Sigma-Aldrich) for 48h with the appropriate doxycycline concentration.

To establish doxycycline inducible system, for RPB1/POLR2A HeLa T-REx (ThermoFisher) were transfected with the Invitrogen Flp-In target site vector, pFRT/lacZeo (ThermoFisher), to introduce an integrated FRT site and generate the host cell line HeLa T-REx Flp-in. Transfected cells were kept under selection until the formation of colonies, with single colonies picked. HeLa T-REx Flp-In host cell line was co-transfected with pFRT-TO-RPB1-His WT^16^, containing 6-His C-terminal tagged RPB1/POLR2A coding and a Flp recombinase expression plasmid, pOG44 (Thermo Fisher), or pOG44 alone, which is the control for the RPB1/PROL2A overexpressing system. For RPB3 and RPB4, HeLa T-REx were transfected with pT-Rex-DEST31 plasmids (Source Bioscience) containing either Empty vector (EV), POLR2C or POLR2D coding region 6xHis N-terminal tagged. All cells were transfected with FuGENE HD Transfection Reagent (Promega) according to manufacturer instructions. Single subunits expression levels were assessed by quantitative PCR (qPCR) and whole cell extract western blot (WB).

### RNA extraction, reverse transcription and Quantitative (Real time) PCR

Cells from a 6-well plate were collected by trypsinization, pelleted by centrifugation, and processed for RNA extraction with RNeasy kit (Qiagen) according to manufacturer’s instructions. 1 μg of RNA was reverse transcribed with random hexamers using the SuperScript III reverse transcriptase kit (Invitrogen), according to manufacturer’s instructions. For Real time PCR (RT-PCR) SYBR Green super mix (Bioline) was used according to manufacturers’ instructions. ACTB was used as control loading gene.

### Transient transcription sequencing (TT_chem_-seq) and library preparation

Nascent RNA was labelled with 1 mM 4-thiouridine (4SU) (Glentham Life Sciences) for 15’ at 37C, and labelling was stopped by removing the culture media and adding 1 mL of TRIzol (Thermo Fisher). Followingly, RNA was extracted according to TRIzol manufacturer’s instruction and samples prepared for TT_chem_-seq as described in Gregersen *et al.* 2020^17^, with 4-thiouracil (4TU)-labelled yeast RNA spiked in for normalization^17^. Library preparation was done with the NEBNext Ultra II Directional RNA Library Prep Kit for Illumina (NEBNext) according to the manufacturer’s instructions. Given the size distribution of the RNA, the protocol followed was the one for “degraded samples”. Libraries were indexed with the 96 Unique dual index primer pairs (NEBNext), quantified, normalized and pooled, and sequenced on Illumina NextSeq system. Each TT_chem_-seq experiment was repeated in duplicate.

### Chromatin immuno-precipitation and library preparation

For chromatin immuno-precipitation (ChIP) 2×15cm dishes of cells were crosslinked with 1% formaldehyde final concentration for 10’ on rotation. Crosslinking was quenched by adding glycine 125 mM final concentration for 5’ on rotation. Cells where then pelleted by centrifugation and chromatin preparation as well as immuno-precipitation of the target proteins were done as previously described^18^. 15 μL of Protein A Dynabeads (Invitrogen) were incubated for 1 h at RT with 1 μg of the corresponding antibody: rabbit anti-Phospho-Histone H2A.X(Ser139) (Abcam); rabbit anti-Phospho-Histone H2A.X (Merck-Millipore); mouse anti Ser5-RPB1 (Cell Signalling); mouse anti NTD-RPB1 (Cell Signalling); rabbit anti Ser2-RPB1 (Abcam). Following all washes, samples were diluted in TE buffer and treated with 1 μL of 20 mg/mL RNAse A (Applichem) for 1 h and 3 μL of Proteinase K (Qiagen) for 2 h at 40C. DNA was purified using a PCR Mini Elute purification Kit (Qiagen) according to manufacturer’s instructions. DNA was quantified with Qubit dsDNAHS Assay Kit (Thermo Fisher Scientific) using a Qubit 3.0 Fluorometer (Thermo Fisher Scientific). Library was prepared with the NEBNext Ultra II DNA Library Prep Kit for Illumina (NEBNext) according to the manufacturer’s instructions. Libraries were quantified, normalized and pooled, and sequenced on Illumina NextSeq system. Each ChIP-Seq experiment was repeated in two biological repeats.

### Bioinformatics analysis

The online platform Galaxy (https://usegalaxy.org^19^) was used for sequencing quality checks using FastQC and mapping of samples. For the TT_chem_-seq samples, reads were aligned using STARRNA^20^ to the hg38 human genome and to the sacCer3 genome for the yeast spike normalization, using existing gene annotation information. The Gene annotation files (GTFs) were downloaded from Ensemble release 110 with reads aligned against each reference genome. The total number of unique mapped reads of the yeast spike in, in the mapped BAM file, was used to calculate a scale factor for the different samples. In each replicate the WT empty vector control (uninduced) sample was used as reference normalizing factor. Strand specific bedgraph files were then created using the SAM tools^21^ to split BAM file in + and – strands and using the advance settings to include the scale factor previously calculated. Read coverage profiles were generated using the EaSeq v 1.101, with the function “Average” around the genetic feature indicated in the figure legends^22^. TT_chem_-seq levels were obtained using the function “Quantify” in EaSeq, from start to end of the gene of all annotated transcribed genes, or else as indicated in figure legends.

For the ChIP-Seq samples, reads were aligned to the hg38 genome using Bowtie2^23^ v.2.3.4.2 on the online platform Galaxy^19^. BAM files from single lanes were merged using SAMtools Merge^21^. Read coverage profiles were generated using the EaSeq v1.101, with the function “Average” around 120% relative to the middle of the gene^22^. ChIP-Seq levels were obtained using the function “Quantify” in EaSeq, from start to end of the gene of all annotated transcribed genes, or as indicated in figure legends. The relative level of gH2AX/H2AX was calculated over the gene or the genomic region. Single genes snapshots were captured on IGV^24^.

### Protein extraction and cell fractionation

For whole cell extract (WCE) the pellet was lysed in RIPA Buffer (10mM Tris pH 8.0, 1mM EDTA, 0.5mM EGTA, 1% Triton X-100, 0.1% Sodium Deoxycholate, 0.1% SDS, 150mM NaCl) supplemented with protease inhibitors (Thermofisher) on ICE for 20’. The WCE was sonicated using the Bioruptor (Diagenode) and samples cleared by centrifugation at 4°C 13000xg for 10’. For cell fractionation, cells collected from a T75 flask were lysed in 200 μL of Cytosolic Buffer (0.15 % NP40 Alt (V/V), 10mM Tris pH 7.0, 150 mM NaCl) for 5’ on ice, using a P1000 with cut tip. Lysate was layered on 500 μL Sucrose Buffer (10 mM Tris pH 7.0, 150 mM NaCl, 25 % Sucrose) and centrifuged at 16000xg for 10’ at 4°C. Supernatant was collected as fraction 1 (cytoplasmatic fraction). Nuclei were washed twice with 800 μL Wash Buffer (1 mM EDTA, 0.1 % (V/V) triton in PBS), each time spin down at 1150xg for 1’ at 4°C. Then, nuclei were resuspended in 200 μL Glycerol Buffer (20 mM Tris pH 8.0, 75 mM NaCl, 0.5 mM EDTA, 50 % glycerol (V/V), 0.85 mM DTT) and lysed in 200 μL Nuclear Lysis Buffer (1 % NP40 Alt (V/V), 20 mM HEPES pH 7.5, 300 mM NaCl, 1 M Urea, 0.2 mM EDTA, 1 mM DTT) vortexing on high 5 seconds and then on ice for 5’. Lysates were centrifuged at 18500xg 4°C for 2’ and supernatant collected as fraction 2 (nuclear fraction). Pellet was washed once in Nuclear Lysis Buffer and once in PBS and then resuspended in 100 μL PBS supplemented with 1.5 mM MgCl2 and 1 μL of Benzonase (5KU; Sigma), and left 1 h at room temperature (RT) or until digested on ice. Digested material was collected as fraction 3 (Chromatin fraction) after a brief spin. All buffers were supplemented with 1 % protease and phosphatase inhibitors, freshly added. For SDS-PAGE, gels were prepared at either 8 % or 12 % (Acrylamide: Bis-acrylamide) run in Running Buffer (Tris Base, Glycine, 20% SDS and transferred using Transfer Buffer (Running Buffer, 20% Methanol) for 1 h and 30’ on Nitrocellulose membrane (BioRad). Membranes were blocked in 5 % milk ((W/V) in TBS (Tris, NaCl) Tween 0.1 % (V/V) (TBST) for 1 h at RT and incubated with the desired antibodies, diluted 1:1000, in 1 % milk in TBST overnight at 4°C. Membranes were washed three times for 5’ each in TBST and incubated with HRP conjugated secondary antibody in 1 % Milk in TBST for 1 h at RT. Before developing, membranes where again washed three times in TBST. Membranes were incubated for 2’ at RT in the dark with ECL (BioRad) and then visualized using the Chemidoc (Bio-Rad) or by using films (Amersham) in the dark room.

### Immunofluorescences and image processing

Cells were seeded on poly-lysine coated slides in 6-well plates and at the appropriate timepoint fixed and permeabilized for 20’ in PTEMF buffer (20 mM PIPES pH 6.8, 10 mM EGTA, 0.2 % Triton X-100 (V/V), 1 mM MgCl2, 4 % formaldehyde (V/V)) at RT and then washed again in PBS for three times, 5’ each. Fixed samples were stored at 4°C until staining. Before antibody staining, cells were blocked for 1 h at room temperature with 10 % FBS/PBS (V/V; Sigma Aldrich), washed in PBS and incubated with 1:500 diluted rabbit anti-53BP1 (Abcam), 1:1000 rabbit anti-Phospho-Histone H2A.X(Ser139) (Abcam) in 1% FBS/PBS for 1h at RT or at 4°C overnight. Cells were then incubated with 1:1000 diluted Alexa Fluor488-conjugated goat anti-rabbit antibody or Alexa Fluor594-conjugated goat anti-rabbit or Alexa Fluor488-conjugated goat anti-mouse (Thermo Fisher Scientific) in 1 % FBS/PBS for 1 h at RT. After being washed in PBS, coverslips were mounted on slides with mounting medium with DAPI (GeneTex). For EU pulse labelling and Click-iT reaction cells were grown in the same conditions as for the immunostaining but pulsed with 50 mM ethyl-uridine (Thermofisher) (EU) for 1 h to label nascent RNA before being fixed and permeabilized in PTEMF buffer. Fixed cells were then processed according to the Click-iT RNA Alexa Fluor 594 Imaging Kit (Invitrogen) manufacturer’s instructions, before either DAPI mounting or primary antibody staining. All images were acquired with Nikon Eclipse E600 microscope. 53BP1 foci images were acquired with 40X objective, EU click-iT and γH2AX images with 60X oil objective. For each sample > 100 nuclei were counted and for a set of samples, the same exposure settings were used. 53BP1 and γH2AX foci, or γH2AX and EU intensity levels were counted/analyzed with CellProfiler 4.0.3^25^. All images were firstly “colored to gray”. Nuclei were identified with “identifyAsPrimaryObjects” module. Nuclei were used to mask foci image with “MaskImage” module, and foci were identified with “IdentifyPrimaryObjects” module Intensity was measured with “MeasureObjectIntensity” module. 53BP1 bodies and mitotic errors (anaphase bridges and lagging chromosomes) were counted manually.

### Cell growth analysis

1000 cells were seeded in a 96 well plate in 50 μL of DMEM in quadruplicate. After 24 h, cells were treated with DMSO, JQ1 500nM or THZ1 1uM, with a top up of media of 50 μL. After another 24 h the first plate was with PBS and stained with 4 % crystal violet (V/V; Sigma Aldrich) in 20 % Methanol (V/V) for at least 30’; excess staining was removed and left to air dry. 10 % (v/v) acetic acid glacial were used to dissolve crystal violet and absorbance was read at 595nm with a plate reader (EnSpire multimode plate reader, PerkinElmer).

## Results

### Upregulations of single subunits of the RNAPII correlates with poor cancer patients’ outcome and increased genome instability

Previous work identified across many cancers multiple deregulations of RNAPII complex subunits, like mutations, copy number alterations and changes in gene expression levels (reviewed in ^7^). However, in many of these studies only the impact of one kind of deregulation was analyzed, and/or any effect of the RNAPII subunit was assessed in the context of the specific cancer type, and not compared across different cancer types or datasets. Therefore, to determine which deregulations are associated with altered patients’ survival, we performed a systematic analysis of all RNAPII subunits deregulations across all TCGA studies available through cBioPortal^14^, to identify the deregulations with the biggest impact. This analysis determined that deregulations of single subunits are frequently found, with 21% of all cancer samples in the TCGA studies presenting a deregulation of at least one subunit, with genomic amplifications the most frequent events (Fig 1A, S1A). Importantly, subunits’ deregulations correlated with reduced overall survival (Fig 1B). When patients were clustered according to the deregulation subtype in “mutations”, “amplifications” or “homozygous deletions”, it was specifically genomic amplifications of any of the 12 subunits that was associated with poor survival (Fig 1B). At the same time, mutations had no effect on survival and patients with genomic homozygous deletions presented a slightly improved survival (Fig 1B). Therefore, we further investigated the impact of gain of functions deregulations of the subunits, combining genomic amplifications with transcriptional upregulations. This allowed us to identify seven separate cancers, where upregulations of the subunits correlated with reduced survival (Fig 1C, S1B). Intriguingly, this was almost invariably due to upregulations of any of the 4 largest subunits clustered together (RPB1-4), separated from the upregulations of the other 7 subunits (RPB5-12) (Fig 1C). This indicated that perhaps upregulations of the largest subunits of the complex could have a bigger impact on patients’ survival than the remaining subunits. To determine whether upregulation of a subunit was associated with increased expression levels of the whole RNAPII complex, we analyzed how the expression levels of RPB1-4 correlated with each other, finding that these were very variable across the characterized cancers, and generally not very high (Fig 1C). To further address this point we determined the correlations between mRNA and protein expression levels of all subunits across hundreds of cancer cell lines using DepMap^26^ (https://depmap.org/portal/), finding again overall poor correlations across all subunits (Fig S1C). This would indicate that in a specific cancer patient, we would be observing higher levels of the single subunit rather than upregulation of the whole complex. Next, we determined whether upregulations of single subunits were associated with specific cancer driver mutations, and whether this was the reason for the correlation with reduced survival. To do this, we analyzed the co-occurrence of subunits upregulations with deregulations with any of the five major cancer drivers specific to the cancer type according to IntOGen^15^ (https://www.intogen.org/search). This showed upregulations of RPB1-4 co-occurring in several cases together with p53 deregulations (Fig S1D). However, patients with deregulations in p53 alone had similar survival rates to patients with only RPB1-4 upregulations in the cancers analyzed, and it was patients with combined deregulation in p53 and upregulation of RPB1-4 that had significantly worse survival than the others (Fig S1E).

**Figure 1.**
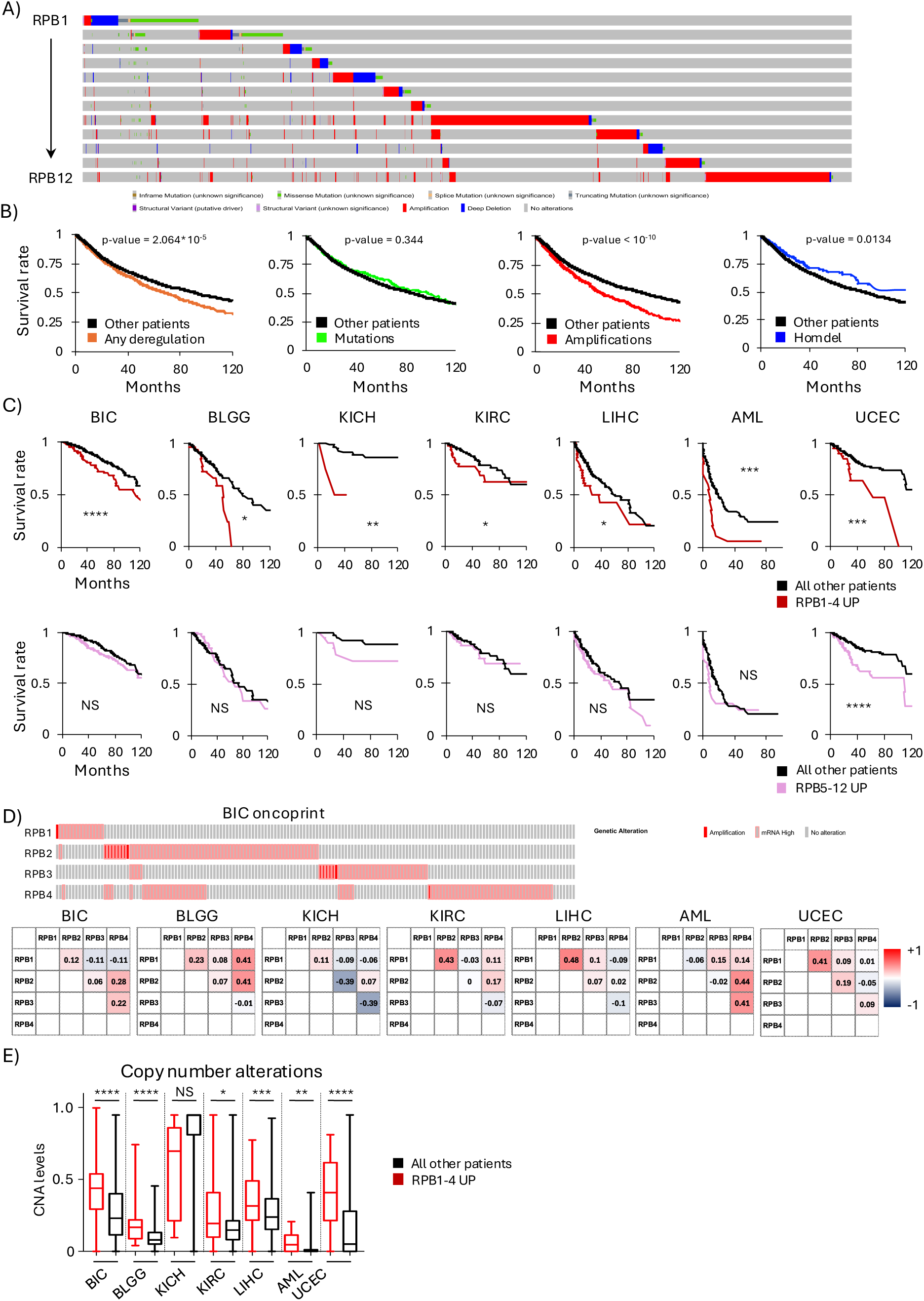
Upregulation of RNAPII subunits correlates with poor cancer patients’ outcome. (A) Oncoprint from cBioPortal of the different types of deregulations for all the subunits of the RNAPII complex. (B) Overall survival for patients with deregulations in any of the RNAPII subunits, either any deregulation or specifically only mutations, amplifications or homozygous deletions (homdel), log-rank Mantel-Cox test. (C) Overall survival in BIC, KICH, KIRC, LIHC, AML and UCEC, for patients with RPB1-4 or RPB5-12 upregulations (amplification + expression > 2 z-score), log-rank Mantel-Cox test. (D) Oncoprint from cBioPortal in BIC specific for upregulations (amplification + expression > 2 z-score) in RPB1-4 and Pearson correlations of the expression levels of all the four subunits in the listed cancers. (E) Comparison of the copy number alterations (CNA) levels in patients with upregulation in RPB1-4 towards all other patients in the listed cancers, box-whiskers plots with line at the median, Mann-Whitney t-test. * => p-value < 0.05, ** => p-value < 0.01, *** => p-value < 0.001, **** => p-value < 0.0001.

We hypothesized therefore that upregulations would lead to non-stochiometric expression of single subunits of the RNAPII. The overexpressed subunit could compete with the RNAPII complex on chromatin for specific transcription factors, potentially affecting RNAPII progression and/or transcription regulation. Deregulated transcription can lead to increased DNA damage in the form of transcription-associated mutagenesis and transcription-associated recombination, consequence of the interference of RNAPII transcription with other processes taking place on chromatin^27, 28^. Hence, we determined whether indeed patients with upregulations of RPB1-4 presented also increased genome instability, finding that overall, they presented higher levels of copy number alterations (CNA) and mutational counts (Fig 1E, S1F). Altogether, this analysis identified that upregulations of the four largest RNAPII subunits correlated with poor cancer patients’ outcome across a range of cancers, presenting increased genome instability, with upregulations appearing not associated with well-established oncogenic deregulations.

### Overexpression of single RNAPII subunits increases DNA damage levels on genes

To test whether overexpression of single subunits affected RNAPII transcription inducing DNA damage, we generated doxycycline-responsive cell lines able to overexpress three RNAPII subunits at the time, RPB1, RPB3 and RPB4, plus empty vector control cell lines. Following doxycycline induction, mRNA levels of the subunits by real-time PCR raised by similar levels in all three cell lines compared to the control cell lines (Fig 2A), visible also at the protein level (Fig S2A). Moreover, the increase in mRNA levels was comparable with the higher expression levels observed in cancer patients with upregulations of the subunits (Fig S2B). First, we analyzed DNA damage markers levels by immunofluorescence. A common DNA damage marker is Ser139-phosphorylation of histone H2A.X (γH2AX), whose phosphorylation is carried out in a redundant manner by different DNA damage sensor kinases^29^. Following overexpression of RPB1/3/4 for 48h we observed in all cases increased global γH2AX, indicating an overall increase in DNA damage levels (Fig 2B). We also quantified γH2AX foci numbers fold change following doxycycline addition compared to the control cell line, to take into account the effect of doxycycline alone on foci formation, finding that only with RPB3 overexpression there was a significant increase in the γH2AX foci number per cell (Fig S2C). Next, we analyzed 53BP1, that forms discreet foci at sites of DNA double strand breaks^30,31^ and bigger structures called bodies evidence of unresolved DNA lesions caused by replication stress which are not correctly repaired before cell division^32^. We found that 53BP1 foci fold changes were significantly increased following overexpression of all subunits (Fig 2C), with bodies also increased but not significantly (Fig S2D). Analysis of other markers of DNA damage like micronuclei and mitotic errors (anaphase bridges and lagging chromosomes), showed that these increased only following overexpression of RPB3 and RPB4 (Fig 2E, Fig S2E). These data indicated that while the overexpression of all subunits induced DNA damage, the specific type of DNA damage could be different between the three subunits, indicating potentially different mechanisms behind the genome instability. Often, deregulated transcription increases DNA damage because of accumulation or persistence of R-loops, three-stranded secondary structures formed by the nascent RNA hybridized with the DNA template strand which can lead to transcription associated mutagenesis and recombination events^27,33,34^. Analysis with the S9.6 antibody which recognizes R-loop structures did not show increased R-loop intensity upon doxycycline induced overexpression of RPB1 and RPB3, and if anything, a 15% average reduction in signal following RPB4 overexpression, indicating that increased R-loops formation is unlikely responsible for the increased genome instability (Fig S2F).

**Figure 2.**
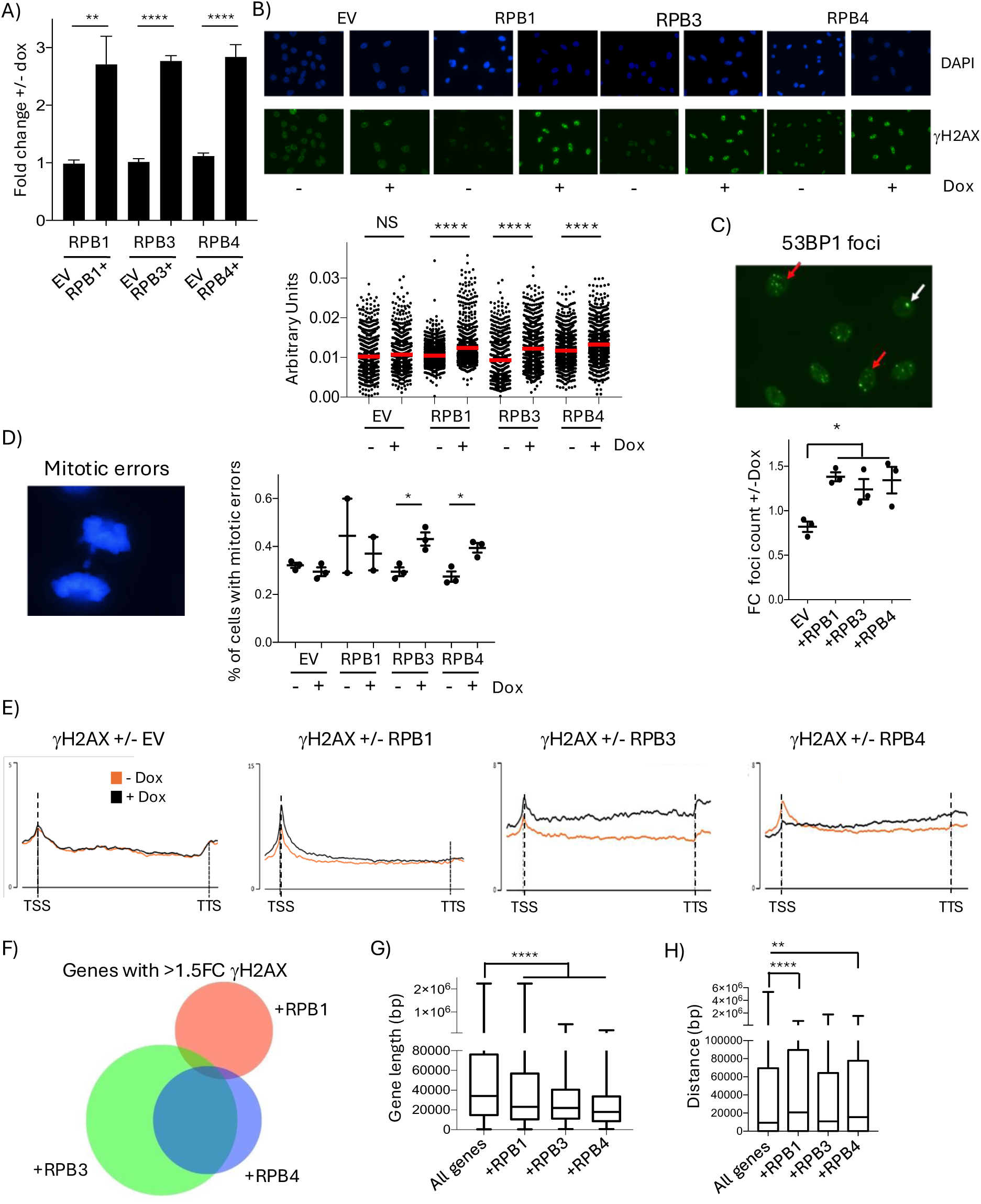
Overexpression of RPB1, RPB3 or RPB4 induces DNA damage in cells. (A) Fold change of RPB1, RPB3 and RPB4 mRNA levels after doxycycline addition in the respective cell lines and the control empty vector (EV) cell line. Average mean +/− SEM, n>=3. (B) Representative images of DAPI and γH2AX immunofluorescence in the listed cell lines +/− doxycycline induced overexpression, and quantification of γH2AX nuclear staining mean intensity, N=3. (C) 53BP1 foci count fold change +/− doxycycline induced overexpression with representative immunofluorescence image, red arrow indicating representative 53BP1 foci while the white arrow indicates 53BP1 bodies, average mean +/− SEM, N=3. (D) Percentage of cells with mitotic errors, counting both anaphase bridges and lagging chromosomes, in the listed cell lines +/− doxycycline treatment, average mean +/− SEM, N=3. (E) Average binned metagene profiles over transcribed genes from TSS to TTS for γH2AX in the listed cell lines +/− doxycycline, a representative profile of the two repeats is shown. (F) Venn diagram showing overlap between genes presenting >1.5 FC in γH2AX levels after RPB1, RPB3 and RPB4 overexpression, averaged for the two repeats. Overlaps between RPB1 and RPB3 (hypergeometric p-value 1.90*10^−23^, under enriched 2.6-fold), overlaps between RPB1 and RPB4 (hypergeometric p-value 8.06*10^−9^, under enriched 2.12-fold), overlap between RPB3 and RPB4 (hypergeometric p-value 1.95*10^−476^, over enriched 4.54-fold). (G) Gene lengths for all transcribed genes and those with >1.5 FC γH2AX following overexpression of the subunits, box-whiskers plots with line at the median, one-way Anova. (H) Distance between transcribed gene and the closest origin of replication mapped in HeLa cells^39^ for all transcribed genes and those with >1.5 FC γH2AX following overexpression of the subunits, box-whiskers plots with line at the median, one-way Anova. * => p-value < 0.05, ** => p-value < 0.01, **** => p-value < 0.0001.

Therefore, to identify and characterize where DNA damage occurred, we performed a chromatin immuno-precipitation followed by sequencing (ChIP-Seq) of γH2AX^18,35^. γH2AX average metagene profiles over transcribed genes showed that overexpression of all three subunits generally increased DNA damage levels, but the actual profiles were different between them (Fig 2E). This agreed with the analysis of the DNA damage markers, highlighting differences between the overexpression of the three subunits induced DNA damage. We found that when we assessed genes with a >1.5 fold-change (FC) increase in γH2AX levels there was a strong significant overlap between genes affected by RPB3 and RPB4 overexpressions, while those affected by RPB1 overexpression where different (Fig 2F). In support of this, we found an overall significant correlation between γH2AX fold changes over all transcribed genes following RPB3 and RPB4 overexpression, but no correlations between γH2AX fold changes following RPB3 or RPB4 overexpression and RPB1 induced DNA damage levels (Fig S2G). On average, genes with more DNA damage were enriched in short genes compared to all transcribed ones (Fig 2G) and were involved in similar biological processes according to their gene ontology (Fig S2H). Altogether, our data indicated that small overexpression of a single subunit was sufficient to induce genome instability in cells, with differences in the DNA damage observed indicating that the mechanisms driving this DNA damage could be potentially diverse between the three subunits. Deregulated transcription can interfere with DNA replication inducing DNA damage, with DNA damage often arising more likely at sites with the most pronounced transcription defect^36,37^, or at the first gene encountered by the replication forks proceeding from replication origins^38^. To test for the latter, we compared the distance from the genes we found accumulating γH2AX following overexpression of the subunits to the nearest replication origin identified in HeLa cells^39^, finding that genes with more DNA damage were not significantly closer to origins than the average transcribed genes (Fig 2H). This was indicating that these genes were perhaps those more transcriptionally affected by the overexpression of the subunit. Hence, we analyzed into details how RNAPII transcription is deregulated following subunits overexpression.

### Overexpression of single subunits alters nascent transcription activity

To analyze RNAPII nascent transcription activity, we performed a transient transcription sequencing, TT_chem_-seq^17^. Cells were pulsed with 4-thiouridine (4SU) for 15 minutes to label newly synthesized RNA, followed by pull-down of the 4SU-labelled RNA and sequencing^17^. Yeast spike-ins were used for normalization to allow quantification of global changes in nascent transcription levels (Fig 3A). Average metagene TT_chem_-seq profiles following subunits’ overexpression showed that they all increased nascent transcription activity (Fig 3A). Upon closer gene to gene analysis and analysis along different gene regions, it appeared that each subunit had slightly different effects on transcription activity. Overexpression of RPB1 increased TT_chem_-seq levels of over 50% from the transcription start site (TSS) to the gene body and the transcription termination site (TTS)+2kb region (Fig 3B) compared to the uninduced samples. In the case of RPB3, TT_chem_-seq levels increased of 36% around the TSS and 30% in the gene body, while only 22% in the 2kb downstream the TTS (Fig 3C). Finally, RPB4 overexpression resulted in increased TT_chem_-seq signal at the TSS and the gene body of 18% and 14% respectively, and 7.4% downstream of the TTS. Based on these observations, we analyzed transcription progression comparing the TT_chem_-seq levels around the TSS (−50/+300bp) to the rest of the gene body (+300bp-TTS) performing a travel ratio analysis^37^, finding small differences following subunits overexpression, more pronounced following RPB3 overexpression (Fig S3A). This indicated that on average, subunits’ overexpression increased RNAPII activity on genes with potential effects also on the transition rate between initiation and elongation and/or altering elongation rates.

**Figure 3.**
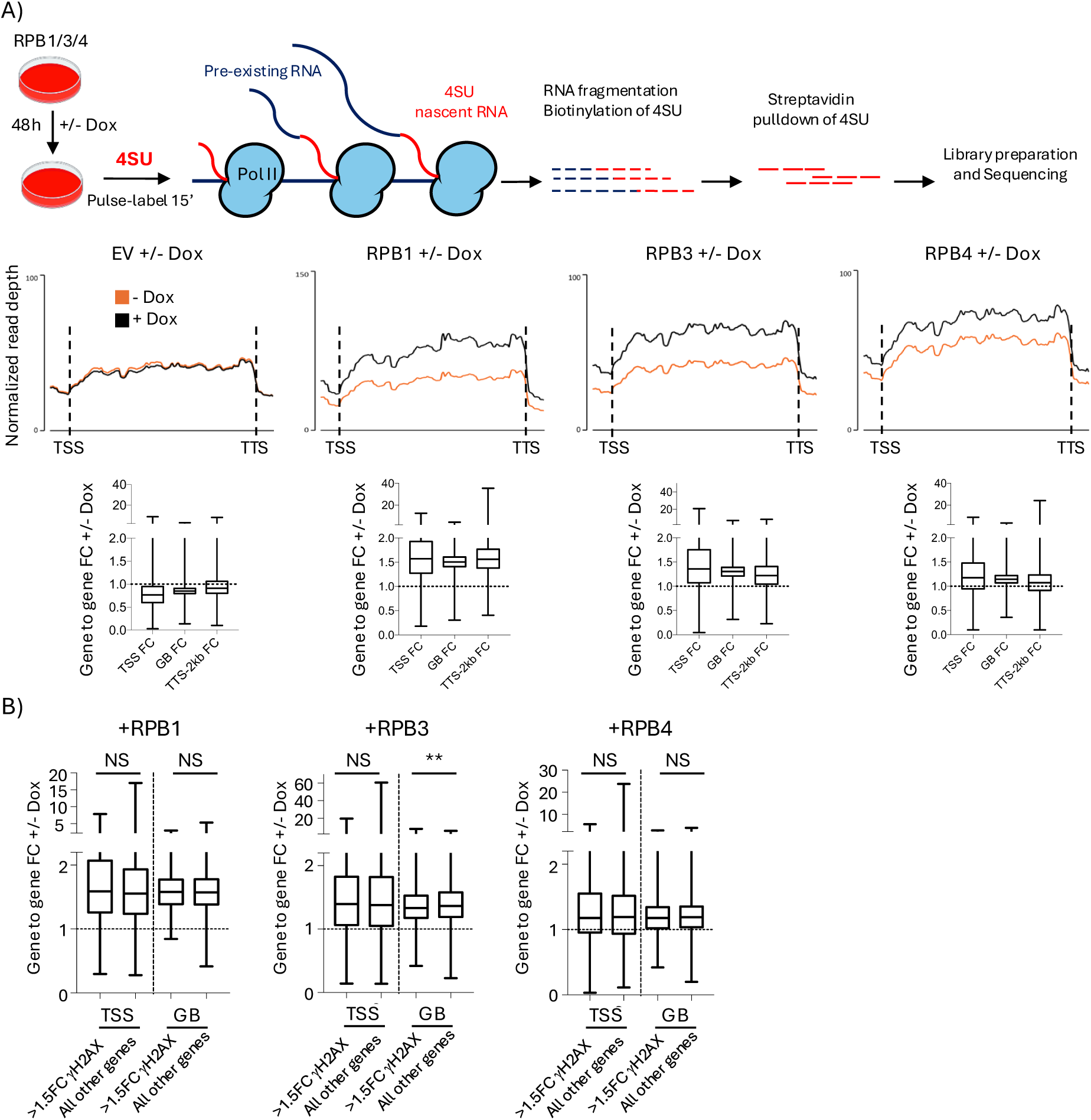
Overexpression of RPB1, RPB3 and RPB4 increases globally transcription activity. (A) Schematic of TT_chem_-seq procedure and average binned metagene profile over transcribed genes from TSS to TTS for strand specific TT_chem_-seq in the listed cell lines +/− doxycycline, a representative profile of the two repeats is shown. Below, normalized quantification of TT_chem_-seq gene to gene fold changes +/− doxycycline over the TSS region (+/− 300 bp), the gene body (GB, TSS+300->TTS), and the TTS+2kb. (B) Gene to gene fold change in TT_chem_-seq fold changes +/− doxycycline addition over the TSS region (+/− 300 bp), the gene body (GB, TSS+300/TTS) for genes with >1.5 FC in γH2AX levels towards all other transcribed genes, box-whiskers plots with line at the median, Mann-Whitney t-test, NS = not significant, ** => p-value < 0.01.

Given that increased transcription activity has been previously linked to increased DNA damage^40,41^, we analyzed whether genes with increased DNA damage were also those more affected by the changes in transcription activity. To do so, we compared the changes in TT_chem_-seq levels on the genes with >1.5 FC in γH2AX levels following the overexpression of the single subunits against all the remaining genes. Intriguingly, there was no difference on how much TT_chem_-seq levels changed around the TSS or within gene bodies on the genes most affected by the DNA damage against all other genes, except for a very small but significant lower FC following RPB3 overexpression in the gene body of genes with more DNA damage (fig 3B). This indicated that there was no direct link between the increased transcription activity on these genes and the increased DNA damage levels. Therefore, we investigated into more details the impact of subunits overexpressions on RNAPII regulation.

### Overexpressions of RPB1, RPB3 and RPB4 alter differently CTD phosphorylation and transcription elongation

To determine the impact of RPB1, RPB3 and RPB4 overexpression on RNAPII transcription, we performed an analysis of RNAPII and its CTD phosphorylated forms Ser2 and Ser5. Ser2-phosphorylation (Ser2-P) and Ser5-phosphorylation (Ser5-P) occur at specific stages of the transcription process and are the best characterized CTD-phosphorylation modifications, important to regulate the interchange of RNAPII-associated factors during transcription^42^. First of all, performing a gene-to-gene fold change of the ChIP-Seq levels upon addition of doxycycline in the EV control cell lines, we found that even 10ng/ml of doxycycline was sufficient to alter RNAPII levels on chromatin and the phosphorylation of the CTD domain (Fig S4A). Therefore, we analyzed the impact of the overexpression of each subunit normalizing for the effect of doxycycline in the empty vector cell lines, as previously done with the DNA damage markers analysis (Fig 2, Fig S2). This analysis found that the overexpression of each subunit affected in a different way RNAPII levels and CTD-phosphorylation. RPB1 overexpression increased globally RNAPII levels on chromatin, on average of 34% in a gene-to-gene comparison (Fig 4A), similarly to what was shown by the TT_chem_-seq data (Fig 3A). At the same time, there was almost no change in Ser2 or Ser5 phosphorylation levels, indicating that when normalized to the total RNAPII levels on genes, this was not properly phosphorylated on Ser2 and Ser5 (Fig 4B, FigS2A). In agreement with the increased TT_chem_-seq signal downstream of the TTS (Fig 3A), we also observed increased total RNAPII levels downstream of specific genes, like highly transcribed ribosomal genes or NABP1 (Fig S4B). This same gene was found similarly affected also upon degron degradation of RPB1^13^, indicating that RPB1 levels, either degron degradation or overexpression, influenced transcription termination. The overexpression of RPB3 presented instead a mixed phenotype. On average we found a global reduction in total RNAPII levels and a profound increase in Ser2-P levels, also visible at the protein level (Fig 4C), with almost no change in Ser5-P. At a more careful analysis, we found that 16.3% of the genes presented a different phenotype, with increased total RNAPII levels and only slightly increased Ser2-P levels instead, and a more pronounced increase in Ser5-P (Fig S4C). Cross-correlating these two groups with the TT_chem_-seq data, we found that they were nearly identical in their TT_chem_-seq level changes either at the TSS or the gene body (Fig S4D), although there was a clear separation in the gene lengths between the two groups (Fig S4E). Altogether, this indicated that following the overexpression of RPB3, a smaller group of genes presented increased RNAPII levels and increased transcription output, while most genes presented increased transcription output with reduced RNAPII occupancy. Finally, overexpression of RPB4 reduced total RNAPII levels on chromatin, slightly increased Ser2 levels and presented a more marked reduction in Ser5-P levels, both in the average metagene and by western blot (Fig 4D). Also in this case, we distinguished two classes of genes, both with reduced Ser5-P levels, 10.7% of genes with increased RNAPII and unaffected Ser2-P, and the rest with decreased RNAPII and increased Ser2-P (Fig S4F). Both groups had nearly identical TT_chem_-seq level changes (Fig S4G) and a clear separation in the gene lengths (Fig S4H). RNAPII ChIP-Seq levels are determined by the number of RNAPII complexes across a region and are profoundly affected by local elongation rates^37,43^. Therefore, the uncoupling between the TT_chem_-seq and RNAPII ChIP-Seq signals would indicate that overexpression of either RPB3 or RPB4 could also affect transcription elongation rates, accelerating RNAPII progression especially along shorter genes.

**Figure 4.**
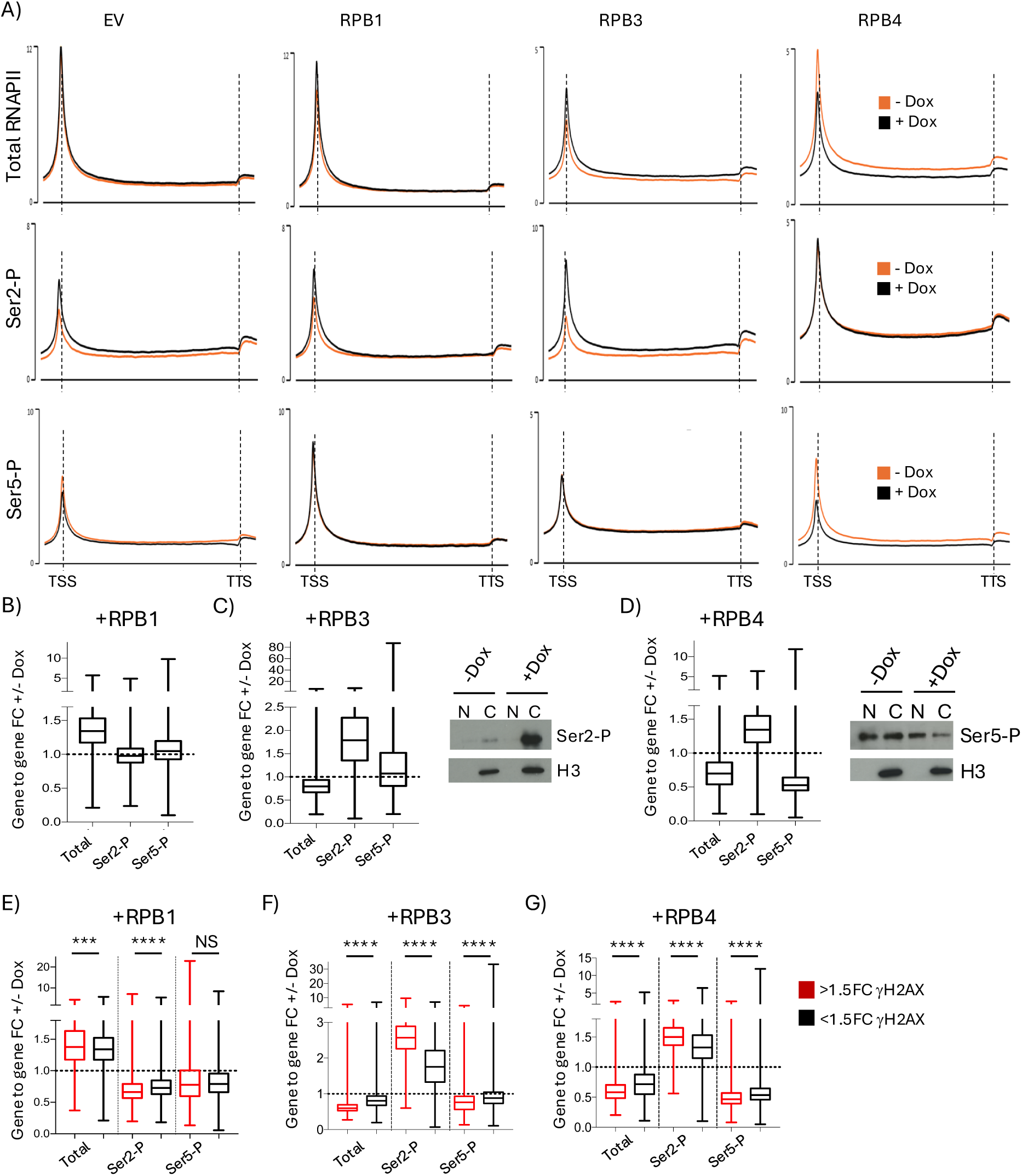
Overexpressions of RPB1, RPB3 and RPB4 alter uniquely RNAPII and CTD phosphorylation levels. (A) Average binned metagene profile over transcribed genes from TSS to TTS for total RNAPII, Ser2-P and Ser5-P in the listed cell lines +/− doxycycline, a representative profile of the two repeats is shown. (B) Gene to gene fold change in total RNAPII, Ser2-P and Ser5-P levels +/− doxycycline-induced overexpression of RPB1, normalized to the impact of doxycycline in the EV cells. (C) As B +/− doxycycline-induced overexpression of RPB3, with western blot for Ser2-P and histone H3 for normalization +/− doxycycline in nucleoplasm (N) and chromatin (C), box-whiskers plots with line at the median, Mann-Whitney t-test. (D) As B +/− doxycycline-induced overexpression of RPB4, with western blot for Ser5-P and histone H3 for normalization +/− doxycycline in nucleoplasm (N) and chromatin (C). (E) Gene to gene fold change in total RNAPII, Ser2-P and Ser5-P levels +/− doxycycline induced overexpression of RPB1 in genes > 1.5 FC in γH2AX following overexpression of the subunits towards the remaining genes, box-whiskers plots with line at the median, Mann-Whitney t-test. (F) As E +/− doxycycline-induced overexpression of RPB3 in genes > 1.5 FC in γH2AX following overexpression of the subunits. (G) As E +/− doxycycline induced overexpression of RPB4 in genes > 1.5 FC in γH2AX following overexpression of the subunits towards the remaining genes, box-whiskers plots with line at the median, Mann-Whitney t-test. NS = not significant, *** => p-value < 0.001, **** => p-value < 0.0001.

Given the complexity of the impact of the subunits’ overexpression on RNAPII regulation and elongation, we analyzed whether the increased genome instability was associated with a specific deregulation. In the case of RPB1 overexpression, we found that genes with a >1.5FC in γH2AX levels presented an even more marked increase in total RNAPII and reduction of Ser2-P, but there was no correlation with the changes in Ser5-P levels (Fig 4E). When we analyzed genes with higher γH2AX levels following RPB3 overexpression we found that these presented overall a more marked reduction in total RNAPII levels, even higher levels of Ser2-P levels and slightly lower Ser5-P levels compared to the rest of the genes (Fig 4F). Indeed, the genes with increased γH2AX levels were almost exclusively within the group of genes with lower total RNAPII levels (Fig S4I). Likewise, genes with increased DNA damage levels following RPB4 overexpression presented overall a more pronounced defect in total RNAPII, Ser2-P and Ser-5 compared to the other genes analyzed (Fig 4G) and were almost exclusively within genes with lower total RNAPII levels (Fig S4J).

Altogether, the ChIP-Seq analysis of RNAPII and RNAPII-CTD phosphorylation patterns showed multiple transcriptional deregulations induced by the overexpression of each single subunit. It also identified how subunits overexpression appeared increasing RNAPII elongation rates, especially over shorter genes. Moreover, it highlighted that not all transcriptional deregulations correlated equally with the increased genome instability.

### Overexpression of single subunits induced deregulated chromatin recruitment of transcription factors, responsible then for the phenotypes observed

Among the transcription-associated factors that interact with the RNAPII complex during transcription, many present domains that recognize CTD phosphorylation status to control their recruitment at specific stages of the transcription process^44,45^. Therefore, considering the widespread changes induced by the overexpressed subunits to the CTD phosphorylation patterns, it was not surprising finding that many transcription factors presented altered chromatin recruitment following subunits overexpression (Fig S5A). More noticeable, BRD4 presented reduced chromatin recruitment following overexpression of RPB3, and reduced interaction with the RNAPII on chromatin by immunoprecipitation (Fig 5A). BRD4, a member of the BET protein family, regulates and activates p-TEFb for Ser2-P on the RNAPII CTD, and it was previously shown that inhibition or down-regulation of BRD4 increased Ser2-P and DNA damage levels^46,47,48^. Thus, we analyzed into more details whether the deregulated chromatin recruitment of BRD4 was responsible for the phenotypes found following RPB3 overexpression. To do this, we overexpressed BRD4 (Fig S5B), finding that this reduced significantly the genome instability and transcription activity induced by RPB3 overexpression, measured by immunofluorescence levels of γH2AX and EU Click-iT incorporation (Fig 5B). Next, we analyzed whether genes with >1.5 FC γH2AX levels following RPB3 overexpression were also those more affected by the loss of BRD4^49^, finding a significant overlap specifically with genes increasing expression following BRD4 loss (Fig 5C). These data therefore would indicate that the overexpression of RPB3 affects the recruitment of BRD4 with the RNAPII, and this can then alter transcription activity and regulation on a subset of specific genes, leading to increased genome instability. Finally, we tested whether this defect in BRD4 recruitment was also sensitizing cells to the BET protein family inhibitor JQ1. In agreement with all the previous data, cells overexpressing RPB3 were specifically sensitive to JQ1 (Fig 5D), further supported by data analysis on the DepMap portal, finding a significant correlation between RPB3 expression levels in 760 cell lines and their sensitivity to JQ1, and a separate BET family inhibitor, I-BET151 (Fig S5C). Altogether, this analysis identified BRD4 as a major factor deregulated by RPB3 overexpression, mostly responsible for driving increased transcription activity and increased genome instability in cells, sensitizing at the same time cells to a JQ1.

**Figure 5.**
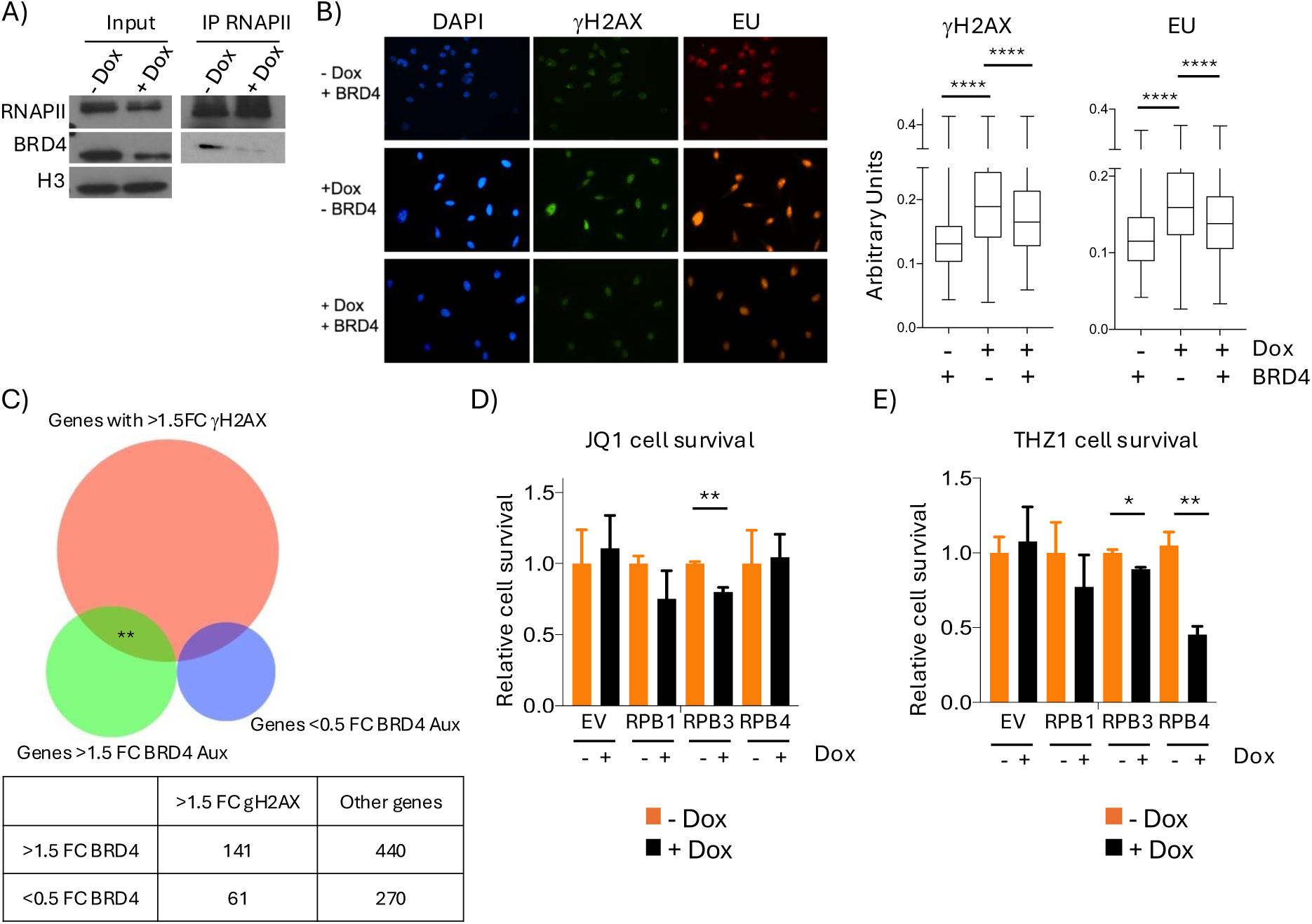
RPB3 overexpression induced deregulation of BRD4 recruitment to RNAPII induced increased DNA damage and transcription activity. (A) Western blot for BRD4, total RNAPII and histone H3 in the chromatin fraction used as input and immunoprecipitation of RNAPII +/− doxycycline addition. (B) Representative images and signal quantification of γH2AX and EU in cells following +/− BRD4 overexpression and +/− doxycycline induced RPB3 overexpression, box-whiskers plots with line at the median, Mann-Whitney t-test. (C) Venn diagram showing the overlap between gene with >1.5 FC γH2AX following RPB3 overexpression and genes >1.5 or <0.5 FC in their transcription following BRD4 Auxin (Aux)-induced degradation from Zheng et al.^49^. (D) Cell survival to JQ1 following doxycycline addition relative to no doxycycline in the listed cell lines, average mean +/− SEM, N=3. (E) As D for THZ1. * => p-value < 0.05, ** => p-value < 0.01.

In the case of RPB4 overexpression, as cells presented reduced Ser5-P (Fig 4F-G), this suggested a potential defect in the recruitment/regulation of the major Ser5-P kinase CDK7, part of the TFIIH complex^42^. Indeed, we found that RPB4 overexpressing cells were sensitive to THZ1, a CDK7 inhibitor, with also RPB3 overexpressing cells presenting a lower level of sensitivity to THZ1 (Fig 5E). To further support a link between the genes presenting higher DNA damage levels following RPB4 overexpression and CDK7 activity, we analyzed how these genes were affected following treatment with THZ1^50^. We found that genes more genome unstable following RPB4 overexpression were downregulated significantly faster compared to other transcribed genes (Fig S5D), indicating that these genes were particularly reliant on CDK7 activity for their transcription. At the same time, cells overexpressing RPB1 that were not presenting only milder defects in Ser2-P or Ser5-P (Fig 4B) were not sensitive to either JQ1 or THZ1 (Fig 5E). Altogether, we have identified how the overexpression of single subunits affected the recruitment of transcription factors to the RNAPII on chromatin, contributing in this way to the deregulated transcription and increased genome instability observed.

### Genes affected by subunits upregulations tend to be also those with increased genome instability in cancers

Our study identified upregulations of RNAPII subunits in cancers correlating with reduced patients’ outcome, determining how single subunits overexpression induced deregulated transcription and genome instability in a model cell line. Therefore, we determined whether there was overlap between the genes with more DNA damage found in our HeLa T-Rex cells after subunits overexpression, and those more frequently genomically altered in the cancers. To do this, we compared how frequently genes with >1.5FC γH2AX levels following subunits overexpression were either copy number altered or mutated in TCGA datasets, compared to control patients without upregulation of the three assessed subunits. First, we analyzed all TCGA datasets together as in Fig 1A-B, comparing patients with an amplified subunit to patients without any subunit affected. The genes with more γH2AX found when we overexpressed subunits presented generally low levels of copy number alterations (CNA) and mutations in control cancer patients without subunits amplifications, indicating that these were not the most genome unstable genes in cancers (Fig 6A). However, in patients with amplification of either RPB1, RPB3 or RPB4, these genes were much more frequently genomic altered, either amplified or deleted, and to a lesser extent accumulated mutations (Fig 6A). Importantly, this was not an indication of increased CNA levels across transcribed genes, as we found for example that in the case of RPB3 and RPB4 genes that presented <0.5FC γH2AX levels following subunits overexpression, were also significantly less CNA compared to genes >1.5FC γH2AX levels in cancers with subunit amplification (Fig S6A). Next, we expanded this analysis to the specific cancer datasets that showed reduced survival following subunits upregulation. In the case of BIC and UCEC, we found strong increases in CNA levels for the genes presenting higher levels of genome instability in cells following overexpression, for each of the subunit analyzed (Fig 6B). In the case of BLGG and KIRP we found increased CNA levels only in cases with RPB4 upregulations, while in LIHC, AML and KICH we found lower levels of genome instability across the selected genes independently of the subunit upregulated (Fig 6B, Fig S6B). These data therefore indicate that at the global pan-cancer level there is a correlation between the genes we found more genome unstable in the cell line model and those more genomically altered in the cancers with subunit’s deregulation, valid also for some of the cancer types initially identified based on the impact on survival. It would also indicate that subunits upregulations, through their deregulated transcription, could directly contribute to the increased genome instability observed across these genes in cancers, linking in this way their upregulations to the reduced survival.

**Figure 6.**
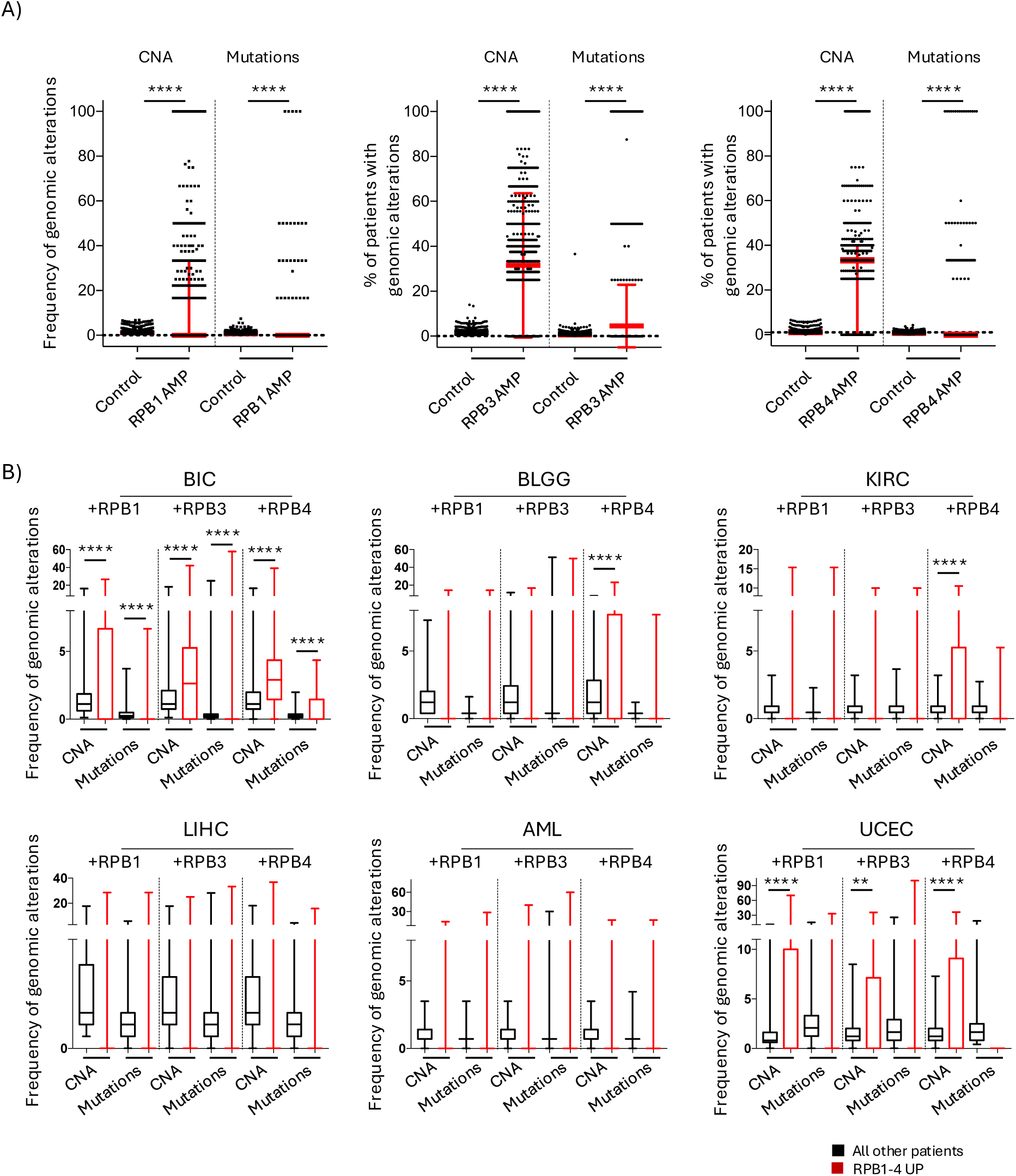
Cancers with upregulations in RPB1, RPB3 or RPB4 present increased genomic alterations across genes found with more DNA damage in the overexpressing cell lines. (A) Percentage of patients with genomic amplification of RPB1, RPB3 or RPB4 in the pancancer TCGA study with CNA or mutations in the genes >1.5 FC γH2AX, compared to percentages of control patients without genomic amplification in the subunits. Minimum to maximum, with line at the median and interquartile range, Mann-Whitney t-test. (B) As (A) but specifically in BIC, BLGG, KIRC, LIHC, AML and UCEC. **** => p-value < 0.0001.

## Discussion

Independently of the specific oncogenic driver, oncogenes will ultimately lead to changes to the transcription program to support their transformative action^51^. Over the years many sequence-specific DNA-binding transcription factors have been found to be highly relevant oncogenes, and more recently this list has been extended also to the general transcription machinery^4^. The RNAPII complex in this sense had been previously found deregulated in many cancers, with deregulations associated with higher cancer proliferation, tumor progression and drug resistance across a broad range of cancer types^7^. Our study has indeed validated that deregulations of RNAPII subunits are frequently found in cancers, with 21% of all cancer samples in the TCGA PanCancer studies presenting at least one genomic alteration in any of the 12 subunits of the complex (Fig 1A). Importantly, by performing a pan-cancer analysis, we found how in particular amplifications of the subunits correlated with reduced cancer patients’ survival, and identified seven cancer types where this was correlating in particular with upregulations in any of the four largest subunits of the complex, RPB1-4 (Fig 1B). While this indicated that upregulations in RPB1-4 correlated more with patients’ reduced outcome than the rest of the complex, it was in no way indicating that it was exclusively RPB1-4 deregulations that were always associated with poor outcome. Another feature shared by cancers with RPB1-4 upregulations was the increased genome instability, in particular CNA (Fig 1E), a typical hallmark of cancer cells^1–3^. Therefore, to determine whether genome instability could be directly induced by RPB1-4 upregulation, we generated doxycycline inducible cell lines able to overexpress one subunit at the time and determined how these overexpressions affected transcription and genome stability. Our data indicated that small overexpression of either RPB1, RPB3 or RPB4, were sufficient to increase DNA damage levels in cells, with subunit-specific differences on how the overexpression impacted on genome stability (Fig 2, Fig S2). A multi-genomic characterization of transcription activity and regulation found that all overexpressions were able to increase transcription activity globally (Fig 3). However, how each subunit overexpression deregulated transcription was subunit specific, and genes with increased genome instability showed changes in transcription activity like those of other transcribed genes. We were able to identify how each subunit overexpression deregulated the CTD phosphorylation pattern of the RNAPII in a unique way, and how genes with increased γH2AX levels presented also the more marked changes to RNAPII and CTD phosphorylation levels by ChIP-Seq (Fig 4). This was particularly noteworthy in the case of RPB3 and RPB4 overexpressions, where we identified distinguished sets of genes with separate changes to total RNAPII, Ser2-P and Ser5-P levels, only one of which linked to the increased DNA damage (Fig 4, Fig S4). This would indicate that increased transcription activity is not on its own sufficient to induce DNA damage in cells, and that not all changes to RNAPII and CTD phosphorylation levels are equally dangerous for genome instability.

RPB1, RPB3 and RPB4 occupy different regions within the RNAPII complex structure and were previously shown important for the interactions between RNAPII and several transcription-associated complexes^10–12,52^. Regarding RPB1, as the largest subunit of the complex that contains also the highly post-translationally modified CTD domain, it could be somehow hypothesised that non-stochiometric expression of RPB1 could affect transcription regulation. Intriguingly, we found that RPB1 overexpression or RPB1 degron degradation^13^ presented a transcription termination defect across the same set of genes, emphasising how RPB1 stoichiometry is essential for transcription termination control. However, our data show that also non-stochiometric expression RPB3 and RPB4 have profound consequences for transcription regulation and genome instability, perhaps even more than the overexpression of RPB1. Cross-referencing our data, we have found instances like the overexpression of RPB1 showing increased RNAPII occupancy and increased TT_chem_-seq signal, indicating overall increased transcription, and others like overexpression of RPB3 and RPB4, where the RNAPII ChIP-Seq occupancy and the TT_chem_-seq signal were going in opposite direction, indicative of a change in elongation rates. Increased transcription elongation rates were previously found following the knock-down of RECQL5, although genome instability was linked to increased RNAPII stalling/pausing (transcription stress) consequence of the uncontrolled elongation^37^. In our case, we found that overexpression of RPB3 and RPB4 gave rise to different deregulations of the CTD, both linked to increased elongation rates in particular across shorter genes. Therefore, we would hypothesise that the increased genome instability is consequence of the CTD deregulation, with its impact on the timely recruitment of transcription factors. The increase in TT_chem_-seq and the reduced RNAPII total ChIP-Seq levels in this sense suggest that stalling/pausing of the RNAPII might not be affected.

Previously, it was shown that RPB3 exists both associated to the RNAPII complex and in a dissociated form, important for the preferential control of 3’-end processing of ribosomal protein genes^53^. When we overexpressed RPB3, we found that ribosomal genes tend on average to present increased total RNAPII levels across the gene, while TT_chem_-seq levels were not particularly affected either on the gene or 2kb downstream of the TTS, indicating that overexpression of RPB3 is not overall affecting ribosomal gene transcription (Fig S6C). Our data showed also how RPB3 was important for the recruitment of BRD4 to the RNAPII on chromatin, regulating CDK9 activity and Ser2-P (Fig 4-5). The significance of RPB3 regulation for this process was represented by how much BRD4 deregulation impacted on genome stability (Fig 4-5). CDK9 is a general transcription factor required globally to regulate RNAPII transcription, however our data indicate that some genes are particularly reliant on its correct regulation, as deregulation of Ser2-P on these genes can lead to increased genome instability. At the same time, we found that 16.3% transcribed genes presented an opposite phenotype following RPB3 overexpression, with increased total RNAPII and very little changes to Ser2-P levels (Fig S4D). These genes were almost completely excluded from the list of genes with more DNA damage (Fig S4E), indicating that transcription deregulations are not equally capable of inducing DNA damage. This if further supported by the lack of correlation between TT_chem_-seq signal and γH2AX changes (Fig 3B). Similarly, in the case of the overexpression of RPB4 we found a global reduction in Ser5-P, with genes with increased DNA damage levels presenting a more pronounced Ser5-P defect (Fig 4), enriched for genes particularly reliant on CDK7 activity for their transcription (Fig S5D). Altogether, this indicates that while transcription deregulation is a major driver or genome instability^27^, not all transcription deregulations will lead to genome instability.

Ser2-P and Ser5-P are two major regulators of RNAPII interactors^45^, and indeed it was not surprising that many of the transcription-associated factors we tested showed increased/decreased chromatin recruitment following subunits overexpression. It is conceivable that among the factors that recognise the CTD phosphorylation status of RNAPII there could be also some important to preserve genome stability in case of transcription-replication conflicts, and that the increased genome instability could also be consequence of deregulated recruitment of such factors to collision sites. Equally, although we have not assessed directly, it is highly likely that also other CTD post-translational modifications might be affected following subunits overexpression, because of crosstalk between CTD modifications and because CDKs can phosphorylate multiple CTD sites^42,54^. At the same time, although our analysis focused on the impact of RPB1, RPB3 and RPB4 because of their more prominent impact on cancer patients’ survival, it is highly likely that overexpression of other subunits may also affect RNAPII transcription inducing DNA damage, with future work needed to determine the impact of the remaining subunits.

### Conclusions and study limitations

Altogether, our study has identified how small overexpression of single subunits of the RNAPII complex can have profound effects on transcription activity, transcription regulation and ultimately genome stability (Fig 7). Several genomic assays have calculated elongation rates in multiple cell lines and conditions^37,55–57^. However, these assays are effective in measuring elongation rates over long genes, and these appear less affected by RPB3 and RPB4 overexpression, making it challenging to apply these assays in these conditions.

**Figure 7.**
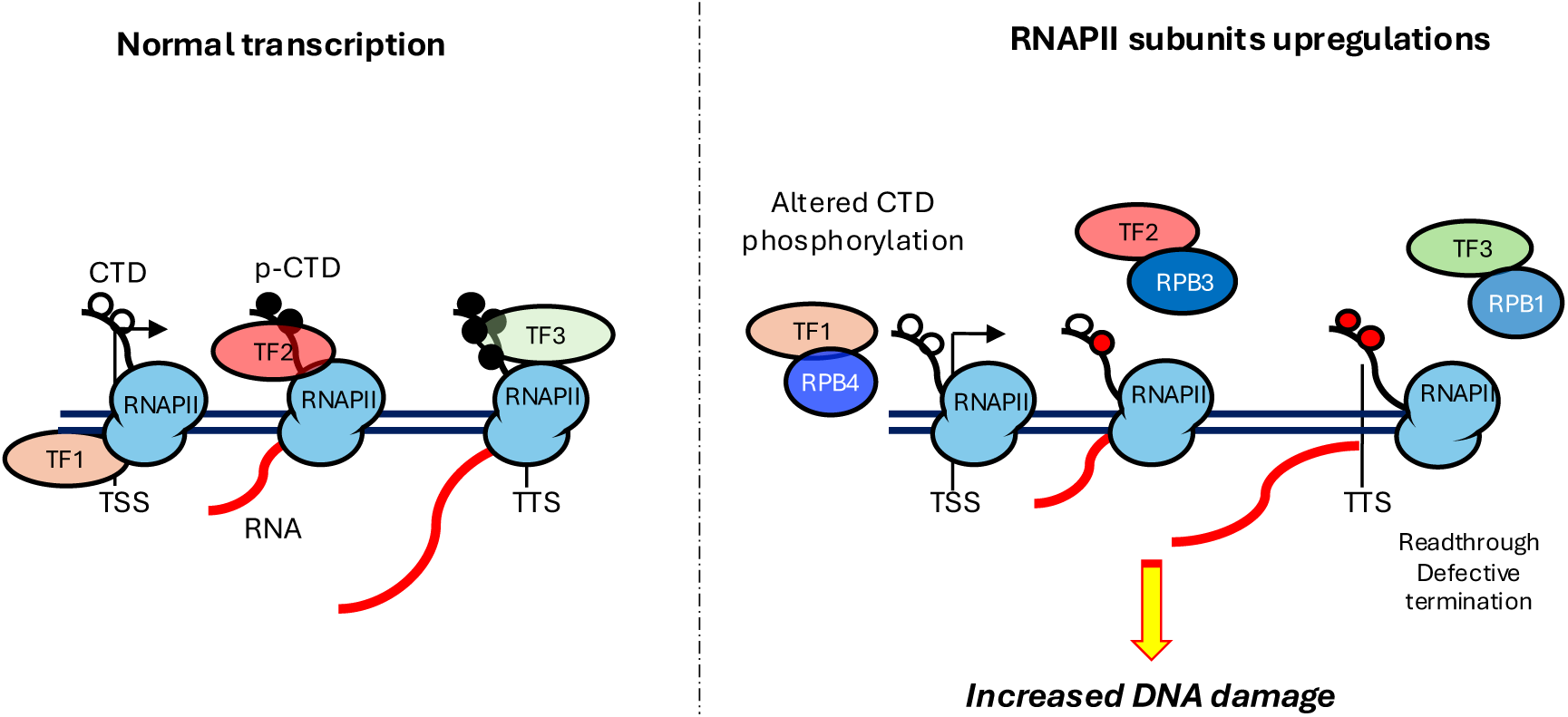
Working model. During transcription progression, the RNAPII interacts with a series of transcription factors, also through the detection of specific post-translational modifications in the CTD associated with that precise transcription stage. However, upregulation of single subunits can interfere with the correct recruitment of transcription factors to the RNAPII complex on chromatin. Among the factors affected there are also the CDKs involved in the CTD phosphorylation, further impacting on the recruitment of transcription factors, inducing deregulated transcription that ultimately leads to increased genome instability.

Intriguingly, we found also a correlation between the genes more affected in our overexpressing cell line (HeLa T-Rex) and those presenting increased genomic alterations in cancer patients baring subunits deregulations. We recognize that this phenotype was only partially recapitulated in the seven cancers we initially identified with reduced patients’ survival correlating with subunits upregulations. Although a well-established cell line, our model cannot recapitulate the phenotypes of all possible cell types, especially as our defects were specifically related to which genes were transcribed in our system. Future work in more appropriate cell lines will be required to determine to what extend subunits upregulations could be a major driver of the genome instability found in these cancers.

## Acknowledgments

This work was supported by the University of Birmingham Fellowship to M.S. and grants from Wellcome Trust (202115/Z/16/Z to M.S.), Royal Society (RG170246, IES\R1\221110 to M.S.), BBSRC (BB/S016155/1 to M.S.), Cancer Research UK (C17422/A25154 to M.S.); work in the lab of L.H.G. is funded by a Hallas-Møller emerging investigator grant from the Novo Nordisk Foundation (NNF20OC0059959), Sapere Aude grant from Independent Research Fund Denmark (0165-00092B), European Research Council Starting grant (ERC-StG-101076758, TranscriptStress) and Center of Excellence (DNRF166) grant from the Danish National Research Foundation (DNFR); work in the lab of A.G. is funded by a Wellcome Trust Investigator Award (215510/Z/19/Z).

## Supplementary figures

**Figure S1:**
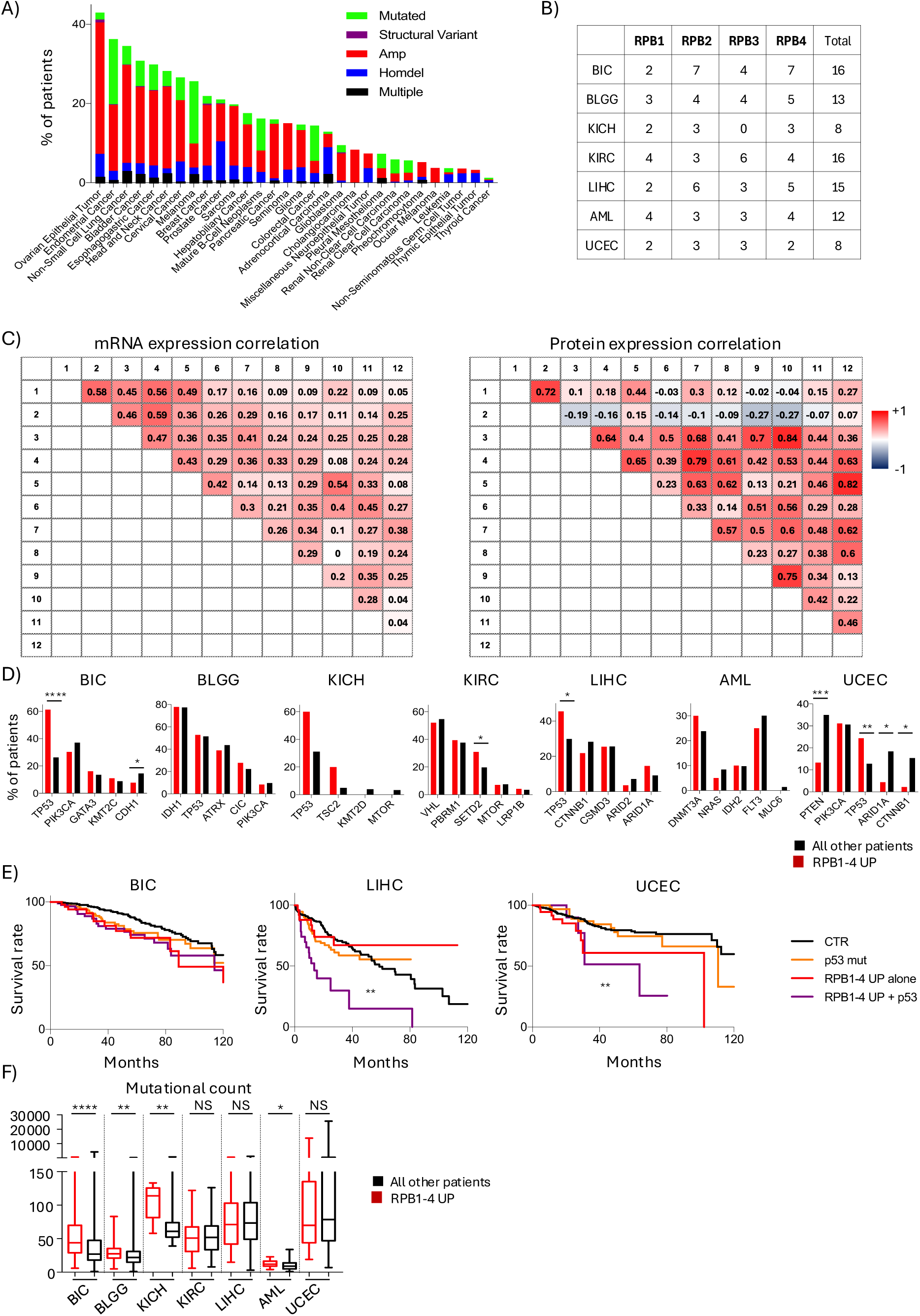
Upregulation of single RNAPII subunits is not associated with a general increase in subunits expression and does not overlap with established oncogenic drivers. (A) Cancer types summary with indicated the percentages of patients with a deregulation in any of the RNAPII subunits, and the breakdown of the specific deregulation. (B) Percentages of patients with upregulation in any of the 4 largest subunits of the complex in the listed cancers. (C) Pearson correlations for mRNA and protein expression from DepMap^26^, with mRNA data from 760 cancer cell lines and protein expression from 375 cancer cell lines. (D) Percentages of patients with RPB1-4 upregulations or all other patients with deregulations in any of the 5 most important cancer driver genes according to IntOGen^15^, expect for KICH with the 4 most important cancer driver genes analyzed because of the size of the cohort. p-value is two-sided Fisher exact test. (E) Survival rate for patients with upregulation of RPB1-4 alone, mutations in p53 alone, patients with upregulation of RPB1-4 and p53 mutations together, and the rest of the patients as controls, log-rank Mantel-Cox test. (F) Number of mutations found in patients with RPB1-4 upregulations against all other patients in the listed cancers, box-whiskers plots with line at the median, Mann-Whitney t-test. * => p-value < 0.05, ** => p-value < 0.01, *** => p-value < 0.001, **** => p-value < 0.0001.

**Figure S2:**
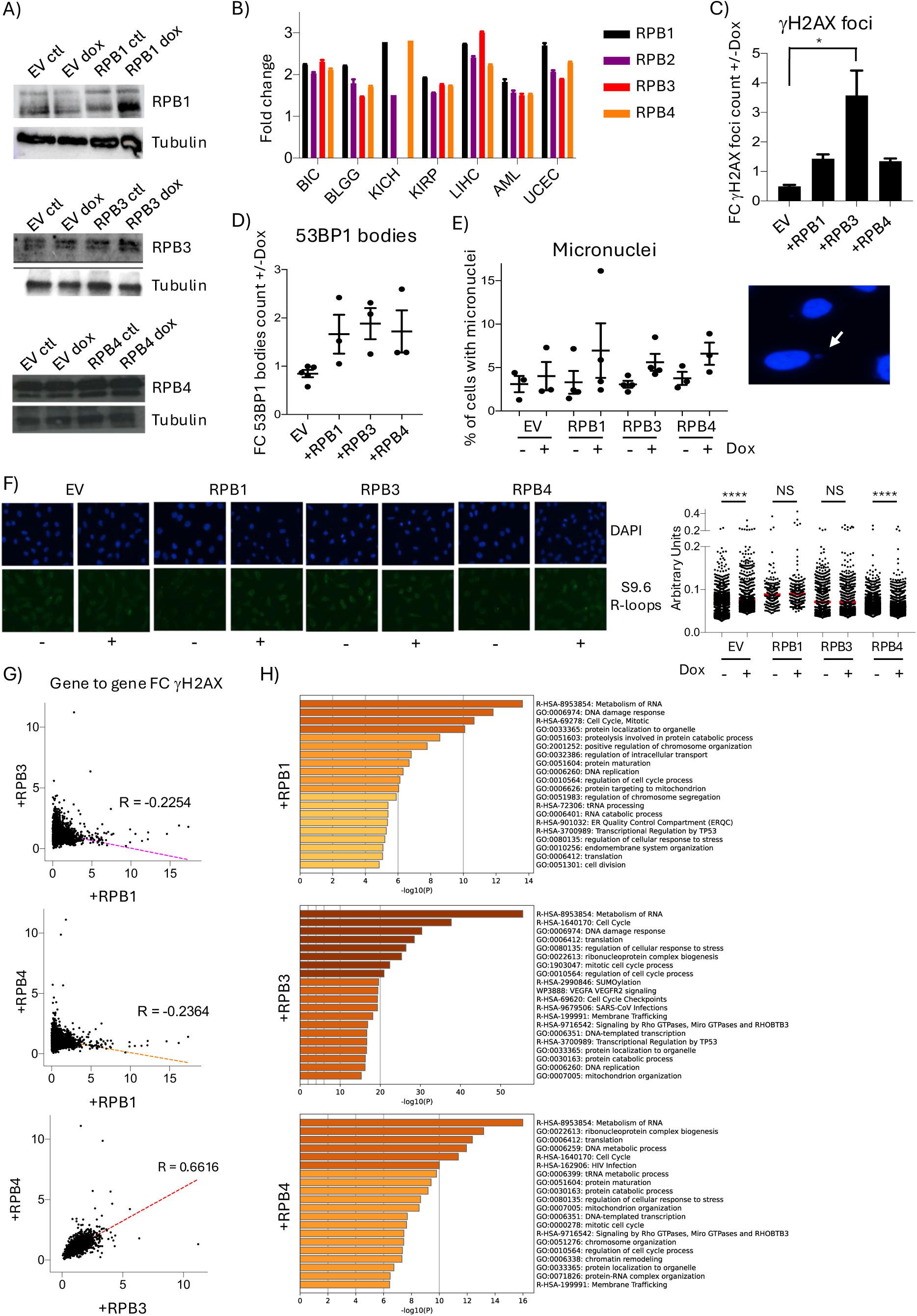
Overexpression of RPB1, RPB3 or RPB4 increases DNA damage levels in cells. (A) Western blot for RPB1, RPB3 and RPB4 in empty vector and doxycycline overexpressing cell line +/− doxycycline, tubulin as loading control. (B) Fold change in average mRNA expression levels in patients with upregulation in the listed RNAPII subunit against all other patients without upregulation. (C) Fold change (FC) in average γH2AX foci count per cell +/− doxycycline. One way Anova, N=3. (D) Fold change (FC) in average 53BP1 bodies count per cell. One way Anova, N=3, not significant. (E) Percentage of cells with micronuclei in the listed cell lines +/− doxycycline, with representative image of micronuclei indicated by the white arrow, N>=3, not significant. (F) Quantification with S9.6 antibody of R-loops levels in the listed cell lines +/− doxycycline, with signal quantification, Mann-Whitney t-test. (G) Pearson correlation for the gene to gene fold change in γH2AX ChIP-Seq levels over transcribed genes +/− doxycycline overexpression of the indicated subunits. (H) Gene ontology analysis with metascape^58^ of genes with >1.5 FC in γH2AX ChIP-Seq levels. NS = not significant, * => p-value < 0.5, **** => p-value < 0.0001.

**Figure S3:**
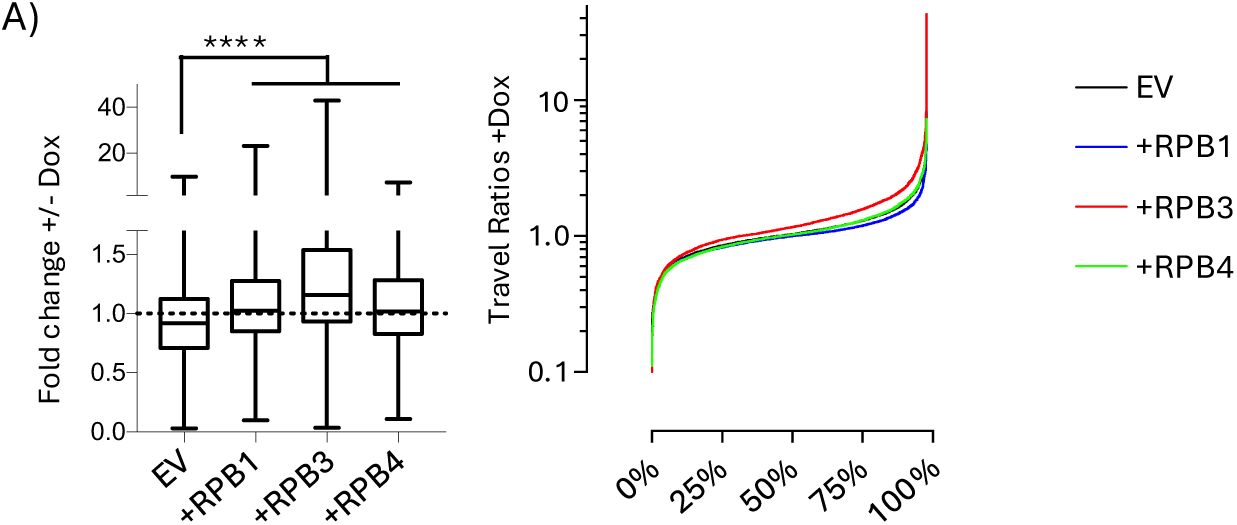
Overexpression of RPB1, RPB3 or RPB4 alters travel ratios. (A) Quantification of the fold changes in gene-to-gene travel ratios +/− doxycycline addition in the listed cell lines, presented as minimum to maximum plot, with line at the median and interquartile range, or arranged in ascending values. One way Anova, **** => p-value < 0.0001.

**Figure S4:**
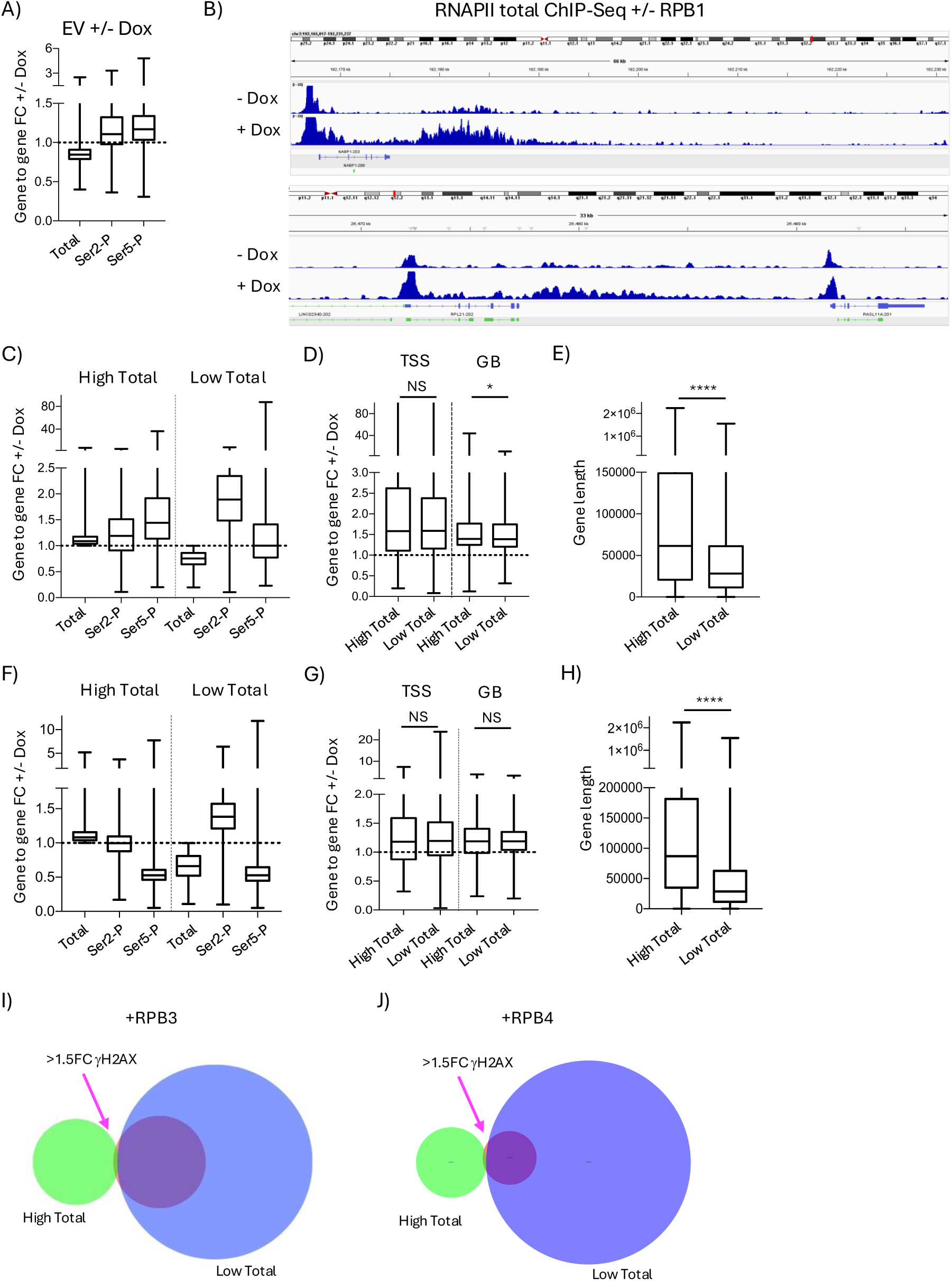
Impact of RPB1, RPB3 or RPB4 overexpression on RNAPII and CTD phosphorylation. (A) Gene to gene fold change in the empty vector (EV) cell line following doxycycline addition for Total RNAPII, Ser2-P and Ser5-P. (B) IGV snapshots of Total RNAPII ChIP-Seq +/− doxycycline addition in the RPB1 overexpressing cell line, highlighting the increased RNAPII levels downstream of the NABP1 and RPL21 genes. (C) Different classes of genes impacted by RPB3 overexpression based the change of total RNAPII levels. (D) TT_chem_-seq gene to gene fold changes at the TSS and gene body (GB) at genes with >1 (High Total) or <1 (Low Total) fold change in total RNAPII ChIP-Seq levels following RPB3 overexpression. (E) Gene length distribution for gene with High Total RNAPII or Low Total RNAPII. (F) As (C) but following overexpression of RPB4. (G) As (D) but following overexpression of RPB4. (I) Venn diagram of the genes with >1.5 FC in γH2AX ChIP-Seq levels and their overlap with the genes >1 (High Total) or <1 (Low Total) total RNAPII fold change ChIP-Seq levels. (J) As (I) but following overexpression of RPB4. Box-whiskers plots with line at the median, Mann-Whitney t-test. NS = not significant, * => p-value < 0.05, **** => p-value < 0.0001.

**Figure S5:**
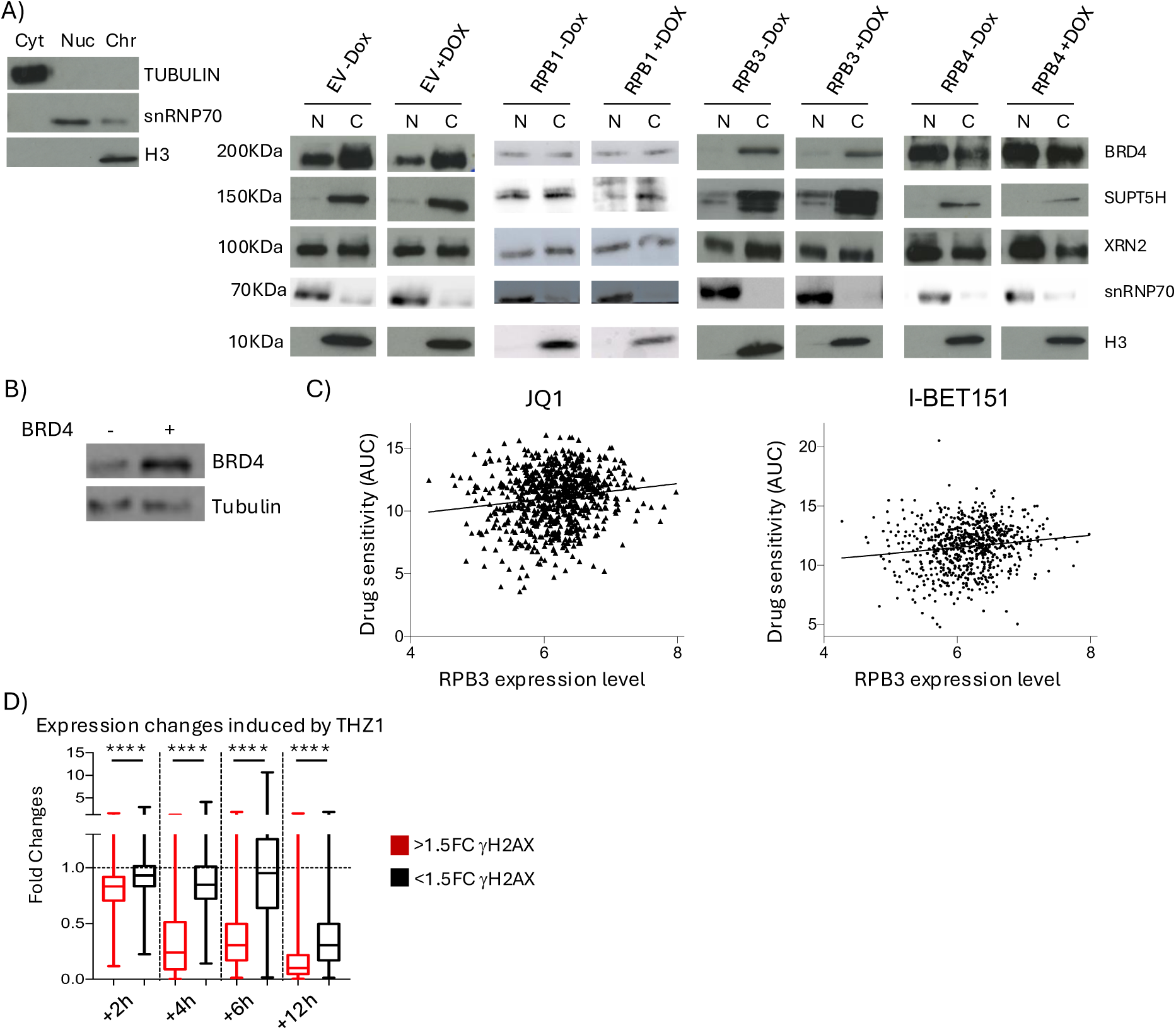
impact of subunits overexpression of defective recruitment of transcription factors to RNAPII. (A) Western blot for SUPT5H, XRN2, BRD4 and histone H3 in the nucleoplasm (N) and chromatin (C) fraction following overexpression of RPB1, RPB3 and RPB4. (B) Western blot for BRD4 following overexpression. Representative western blot shown, N>=2. (C) Scatter plots of RPB3 expression level and cell lines’ drug sensitivity to BET family inhibitors JQ1 (R=0.146, p-value 5.3*10^−5^) and I-BET-762 (R=0.135, p-value 3.1*10^−4^) in 760 cell lines from DepMap. (D) Fold change in expression following treatment with THZ1 after 2/4/6/12 hours for genes with >1.5 FC in γH2AX ChIP-Seq following RPB3 overexpression against all other transcribed genes, box-whiskers plots with line at the median, Mann-Whitney t-test. **** => p-value < 0.0001.

**Figure S6:**
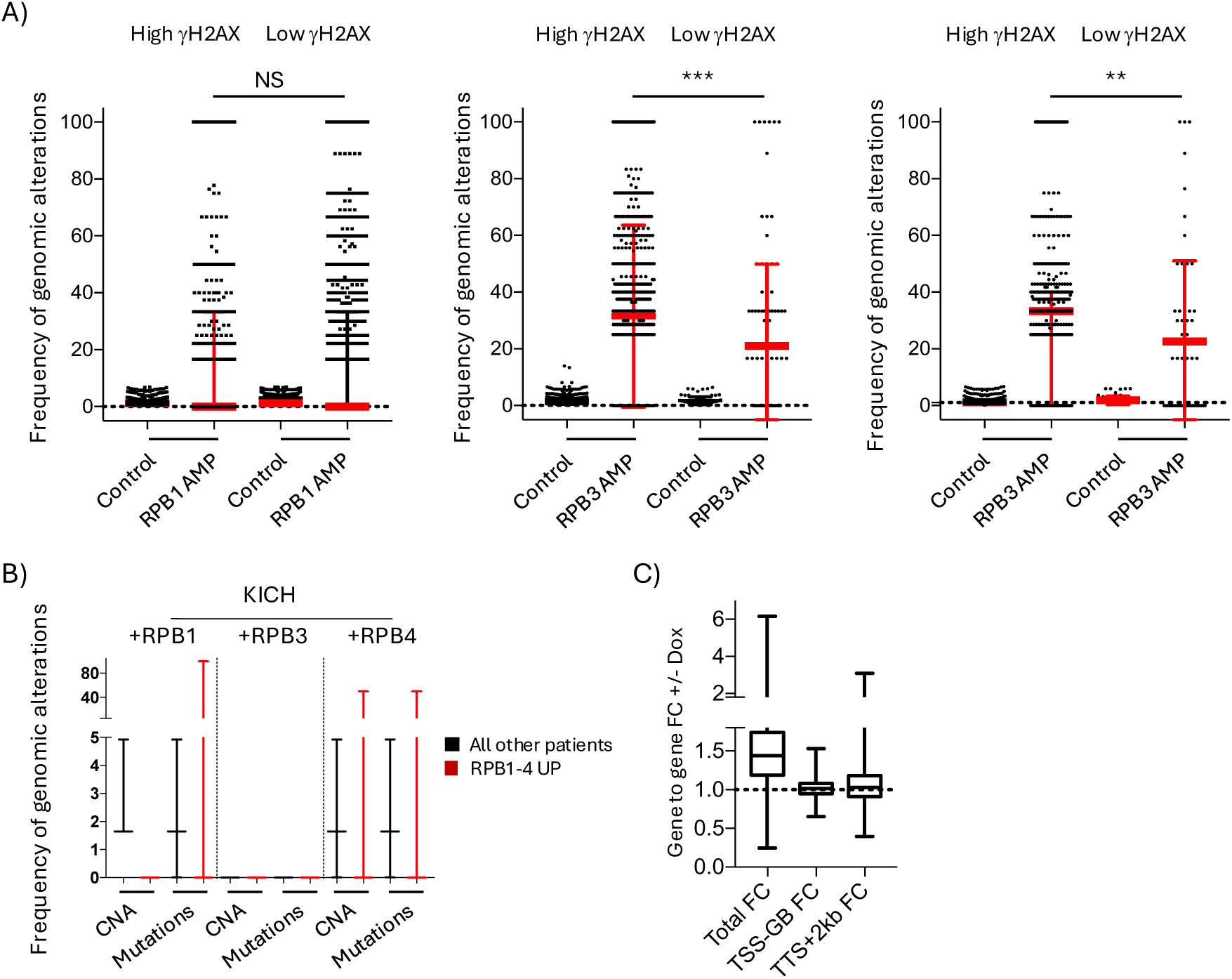
genes with more γH2AX ChIP-Seq signal following subunits’ overexpression are also more frequently genomic altered in cancers with subunits upregulations. (A) Frequency of copy number alterations in genes with >1.5 or <0.5 FC in γH2AX ChIP-Seq levels following overexpression of RPB1, RPB3 or RPB4, in patients with genomic amplification of the RNAPII subunit against control patients without subunits’ amplification. Minimum to maximum, with line at the median and interquartile range, Mann-Whitney t-test. (B) As Fig 6B but specifically in KICH. NS = not significant, **** => p-value < 0.0001.

